# Brain-wide connectivity map of mouse thermosensory cortices

**DOI:** 10.1101/2022.06.29.498101

**Authors:** Phillip Bokiniec, Clarissa J. Whitmire, Tobias M. Leva, James F.A. Poulet

## Abstract

In the thermal system, skin cooling is represented in the primary somatosensory cortex (S1) and the posterior insular cortex (pIC). Whether S1 and pIC are nodes in anatomically separated or overlapping thermal sensorimotor pathways is unclear, as the brain-wide connectivity of the thermal system has not been mapped. We address this using functionally targeted, dual injections of anterograde viruses or retrograde tracers into S1 and pIC. Our data show that inputs to S1 and pIC originate from two non-overlapping populations, supporting the existence of parallel input pathways. While outputs from S1 and pIC were more widespread and share a number of cortical and subcortical regions, within target structures axonal projections were often separable. We observed a high degree of reciprocal connectivity with thalamic and cortical regions from both S1 and pIC, but output to the mid- and hind-brain was unidirectional. Notably, pIC showed exclusive connectivity with regions associated with thermal processing. Together, these data indicate that cutaneous thermal information is routed to the cortex via multiple, parallel streams of information which are forwarded to overlapping downstream regions for the binding of complex somatosensory percepts and integration with ongoing behavior.

## INTRODUCTION

A fundamental feature of mammalian sensory pathways is that the same modality is represented in multiple cortical areas. As the circuitry of different cortical sensory regions is typically studied independently, the neuronal wiring principles of multiple sensory representations is unclear. Different cortical sensory representations could be separate nodes in anatomically segregated, ‘parallel’ neural pathways (Fig. 1a - left). Alternatively, the same presynaptic nuclei could provide copies of sensory information to widespread cortical regions for forwarding to overlapping brain areas in a ‘mixed’ connectivity model (Fig. 1a - right). The thermal system is an ideal model system to address this question as both the primary somatosensory cortex (S1) (Hellon et al., 1973; Tsuboi et al., 1993; Milenkovic et al., 2014) and the posterior insular cortex (pIC) (Penfield and Faulk, 1955; Craig et al., 2000; Beukema et al., 2018; Vestergaard et al., 2022) are involved in thermal processing. Moreover, our recent work has shown that both S1 and pIC have a rich cellular representation of cooling and powerful role on thermal perception (Milenkovic et al., 2014; Vestergaard et al., 2022). However, despite the importance of temperature for somatosensation, there is no comprehensive connectivity map of the mouse thermal system (Bokiniec et al., 2018).

**Figure 1.**
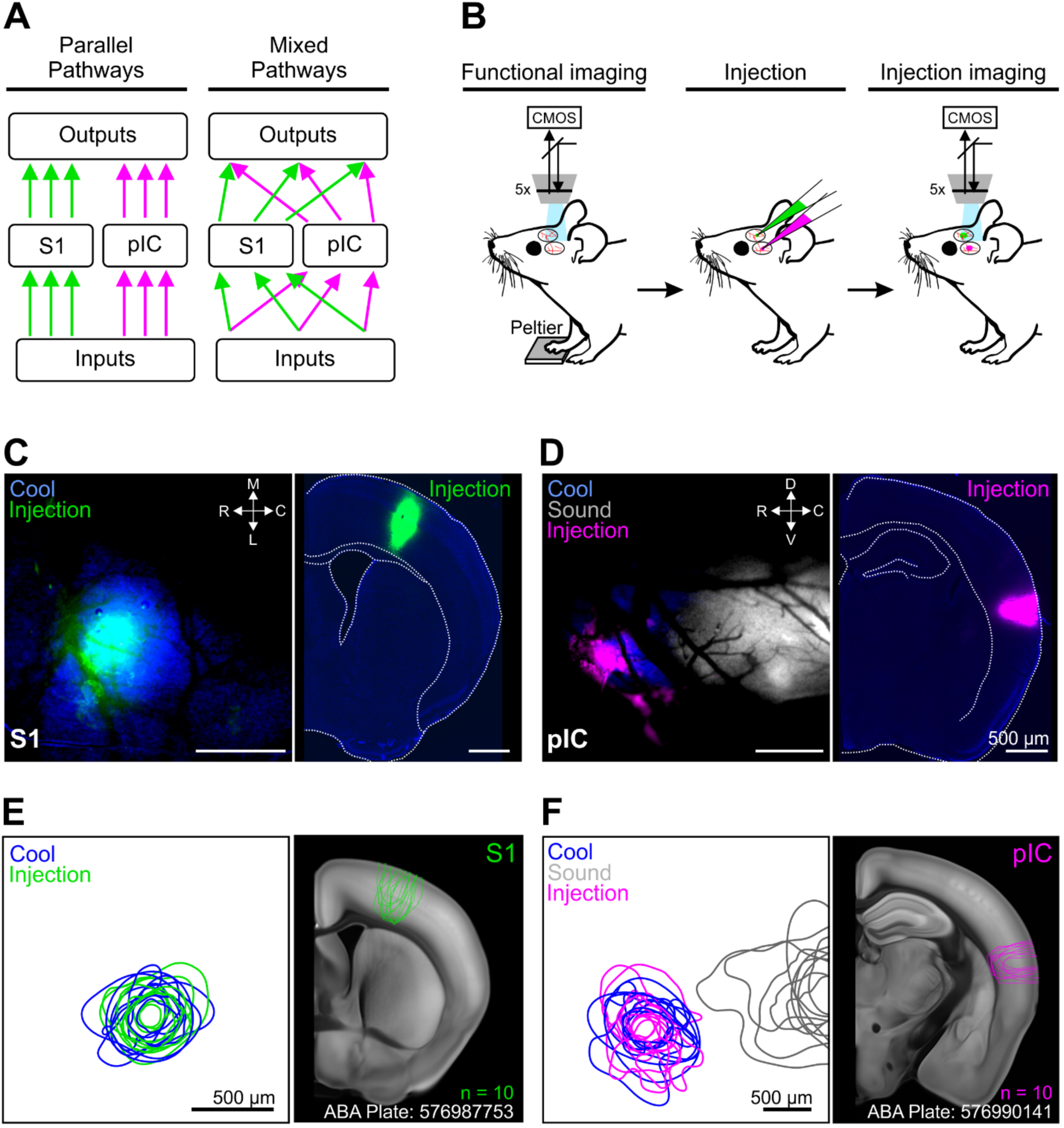
Functional identification of temperature representation in S1 and pIC for anatomical input-output tracing. **A**, Cartoon schematic showing segregated and mixed connectivity motifs. **B**, Schematic representation of the experimental procedure; from left to right, functional identification of the thermal cortical regions using widefield calcium imaging through a cleared skull preparation; injection of different color retrograde tracers or anterograde viruses; imaging to confirm alignment of injection sites to thermal representation. **C**, Example mouse imaging of S1 and corresponding coronal brain slice. Left, overlaid functional response to temperature (blue), with fluorescent tracer (pseudo-coloured green); right, post-hoc brain slice showing infection site. Scale bars 500 µm. **D**, Same as (**C**), but for pIC. **E**, Population injection sites and functional responses over S1. Left shows 80 % contours of the widefield thermal response to cool stimuli (blue) and fluorescence of the tracer (green) (n = 10 mice, 5 retrograde and 5 anterograde injections) aligned to peak temperature response in S1. Right shows outlines of all injection sites localized on coronal brain slice 54 from the Allen Brain Atlas. **F**, Left, same as **E** (left), but for pIC and including response to 8 kHz sound stimulation (gray). Right, same as **E** (right), but coronal brain slice section 70 of the Allen Brain Atlas.

The connectivity of forelimb S1 and IC has been examined independently in prior studies (Guldin and Markowitsch, 1983; Cechetto and Saper, 1987; McDonald and Jackson, 1987; Allen et al., 1991; Shi and Cassell, 1998a, 1998b; Kimura et al., 2010; Maffei et al., 2012; Oh et al., 2014; Zakiewicz et al., 2014; Zingg et al., 2014; Gehrlach et al., 2020). Independent tracing allows a comparison of large-scale wiring differences between two regions, but prohibits a comprehensive examination of subregion and cell-specific connectivity. Moreover, most prior work has used stereotactic targeting for tracer injections. Because blood vessel patterns as well as brain and skull sizes vary from mouse to mouse, it is challenging to determine whether sterotactically-targeted cortical regions correspond to specific sensory representations.

Here we performed functional widefield calcium imaging to target tracer injections to the thermal representations in forelimb S1 and pIC. To allow a direct comparison of inputs and output wiring of S1 and pIC, both regions were injected in the same mouse. We used anterograde adeno-associated viruses (AAV) to trace axonal projections (Viswanathan et al., 2015) or cholera toxin subunit B (CTb) for retrograde tracing of cellular resolution inputs. Brains were then sliced and imaged from hindbrain to frontal cortex, and the brain-wide input and output connectivity from forelimb S1 and pIC were quantified using automated cell counting and axon density estimates. Our study provides a comprehensive whole-brain connectivity map of two major thermal cortical representations, and suggests that there are two independent thermal pathways routed via S1 and pIC.

## MATERIALS AND METHODS

### Mice

All experiments were approved by the Berlin Landesamt für Gesundheit und Soziales (LAGeSo), and carried out in accordance with European animal welfare law. Adult (n = 10), male and female GP4.3 (C57BL/6J-Tg(Thy1-GCaMP6s)GP4.3Dkim/) mice from Jackson Laboratories (JAX#024275, Chen et al., 2013) were used. Mice were housed under 12-hour light/dark cycles and provided with *ad libitum* food and water.

### Surgery

Mice were anaesthetized with isoflurane in oxygen (3-4 % initiation, and 1-1.5 % maintenance, CP-Pharma) and injected with Metamizol for post-operative pain management (200 mg/kg, s.c., Zentiva). Anaesthetized mice were then placed in a nose clamp and eye gel (Vidisic, Bausch + Lomb) was applied to both eyes. Core temperature of mice was maintained by a homoeothermic heating blanket (FHC). The right forepaw was tethered onto a Peltier element (8x8 mm, Digi-Key Electronics) for thermal stimulation of the paw. The left primary somatosensory cortex forelimb (S1) was exposed by removing all the skin on the parietal bone and locating Bregma and Lambda suture landmarks. The left posterior insular cortex (pIC) was exposed by rotating the head ∼30-40 degrees to the right and by displacement of the left temporalis muscle from the temporal bone. The rhinal vein, middle cerebral artery and zygomatic bone were used as anatomical landmarks for pIC (Vestergaard et al., 2022). The skull overlying forelimb S1 and pIC was thinned with a dental drill (head-diameter: 10 mm, Komet Dental) to improve image quality.

### In vivo imaging

Widefield calcium imaging was used to identify thermal locations in S1 and pIC as previously described (Vestergaard et al., 2022). Briefly, images were acquired by a sCMOS camera (Hamamatsu ORCA-Flash4.O LT) via an epifluorescent stereomicroscope (Excitation: 470/40 nm, Emission: 525/50 nm, Leica MZ10 F) equipped with a CoolLED pE-300 LED Microscope illuminator, at a rate of 20 Hz with 35 ms exposure time. Thermal stimuli were delivered to the right forepaw via a feedback-controlled Peltier element stimulator (custom made device, ESYS GmbH Berlin). Cooling stimuli were 10 or 14 °C drop from 32 °C baseline with a duration of 2 s and onset ramp of 20 °C / s. The location of pIC was further confirmed by identifying the surrounding auditory cortex, and, in some cases, the insular auditory field (see: Rodgers et al., 2008; Sawatari et al., 2011; Gogolla et al., 2014) using an 8 kHz, ∼65 dB, 1 s acoustic sound via a loudspeaker (Visaton). Mice received a minimum of 3 stimulation trials to confirm functional responses. Craniotomies (∼1x1 mm) were then performed over the thermally responsive regions in S1 and pIC.

### Tracer and virus injections

Custom written code (Python version 3.7, Python Software Foundation) was used to identify the center point of the widefield response during the imaging session. The center-of-mass was computed from all pixels above 80 % of the peak fluorescence in the trial-averaged responses. Next, glass pipettes (∼ 20 µm diameter) containing cholera toxin subunit b (Ctb), for input mapping, or adeno-associated viruses (AAV), for axonal output mapping, were inserted into the center of the thermal response, normal to the cortical surface. Two 50-75 nL injections (100 nL/min) were made, one at 700 µm and a second at 400 µm depth from the pial surface, using an oil hydraulic manipulator (One-axis oil hydraulic micromanipulator, Narishige). Pipettes were left in place for 5-10 min following each injection and then slowly retracted. S1 was injected with either Ctb Alexa Fluor 647 (Ctb-647, 0.5 % in PBS, Thermo Fisher) or AAV-smFP-myc (pAAV.CAG.GFPsm-myc.WPRE.SV40, 7.17 x 10^11^ vg/ml), and pIC with either Ctb Alexa Fluor 555 (Ctb-555, 0.5 % in PBS, Thermo Fisher) or AAV-smFP-FLAG (pAAV.CAG.Ruby2sm-Flag.WPRE.SV40, 1.58 x 10^12^ vg/ml). To visualize AAV cortical injection sites, AAVs were mixed with a low concentration of Ctb Alexa Fluor 488 (0.05 % v/v, 0.5 % in PBS, Thermo Fisher).

To confirm the location of the injection site, we imaged the fluorescence tracer 10 min post injection while on the imaging setup with the same angle, orientation and field-of-view using either orange light (Excitation: 575/70 nm, Emission: 640/50 nm) or a green LED light at 20 Hz, 35 ms exposure time. A small layer of bone wax was then placed over both craniotomies to prevent tissue damage. The exposed skull was then covered with dental cement (Paladur) and mice placed onto a 37 °C heating blanket and finally returned to their home cage. Drinking water was supplemented with Metamizol (200 mg/kg, Ratiopharm) for post-operative pain management for 2-3 days.

### Histology

Five to seven days after injection of Ctb, or three to four weeks after injection of AAV, mice were anaesthetized with an overdose of ketamine/xylazine (1200 mg/kg ketamine, 500 mg/kg xylazine, i.p., WDT eG and Bayer, respectively) and transcardially perfused with 50 mL ice-cold PBS (0.1 M) followed by 50 mL of ice-cold 4 % PFA. Brains were removed and post-fixed overnight in PFA at 4 ⁰C. Whole brains were cut into coronal sections (50 µm) using a vibrating microtome (Leica VT1000S) and every 4^th^ section was collected. Sections containing Ctb were directly mounted onto glass slides using DAPI Fluromount-G (Southern Biotech) mounting medium.

Sections containing AAVs were stored for further immunohistochemical processing as described previously (Bokiniec et al., 2017). Briefly, free-floating sections were first washed in PBS containing 0.3 % Triton X-100 (3 x 10 min, RT) and then blocked with 5 % normal goat serum in the above wash solution for 60 min at RT. Sections were incubated in primary antibodies (diluted in the blocking solution, see Table 1) against myc (rabbit c-Myc, 1:1000, Sigma-Aldrich, C3956, RRID: AB_439680) and FLAG (mouse-FLAG, 1:1000, Sigma-Aldrich, F1804, RRID: AB_262044) for 48 h at 4 ⁰C. Sections were washed with PBS and then incubated in PBS containing 5 % normal goat serum with fluorescent conjugated secondary antibodies (Alexa Fluor 555-conjugated goat anti-mouse IgG, 1:500, Thermo Fisher, A21422, RRID: AB_2535844, and Alexa Fluor 647-conjugated donkey anti-rabbit IgG, 1:500, Thermo Fisher, A31573, RRID: AB_2536183) overnight at 4 ⁰C. Brain sections were then washed and mounted onto glass slides using DAPI Fluromount-G (Southern Biotech) mounting medium.

Brain sections were visualized with a Zeiss upright microscope (Axio Imager A.2) using the ZEN Imaging software. Images were acquired using a 10x/0.45NA objective. Exposure times for AAV or Ctb were kept the same across mice.

### Histological image processing

#### Atlas registration

Images were first separated by fluorophore, organized sequentially, rotated to the correct orientation, and down sampled (20 % from original) using ZEN Imaging software. Using the ImageJ plugin Fiji (Schindelin et al., 2012), a 1 mm boundary in the rostral-caudal, medial-lateral, and dorsal-ventral axis was masked over the center of the injection sites and excluded from further analysis. All slices were registered to the Allen Brain Atlas Common Coordinate Framework v3 (ABA) using the QUICKNii software package (Puchades et al., 2019). Due to possible section distortion along the dorsal-ventral, rostral-caudal or medial-lateral axes as a consequence of histological processing, images were adjusted using QUICKNii. Sections were contrast adjusted in QUICKNii to allow clear matching of anatomical landmarks from the slice to the atlas. Following complete registration of the sections to the ABA, the corresponding RGB atlas images were exported from the QUICKNii software.

#### Signal detection

Cell soma were identified using a modified version of AIDAhisto (Pallast et al., 2019) that allows interaction with the ABA RGB atlas (MATLAB Version R2018b, The MathWorks Inc.). Images were filtered using the Leung-Malik Filter Bank (Leung and Malik, 2001) to detect non-circular cells with a size between 8 - 10 pixels (corresponding to 20 - 25 µm in the down sampled image). A single threshold for cell detection was determined empirically and was applied to all the datasets. To reduce the identification of false positive cells, the XY cell positions were referenced to a corresponding binarized DAPI nuclei image using the *k*-Nearest-Neighbor classification where *k* = 1, within a radius of 1.5 pixels. Detected cells were then manually checked to their corresponding micrographs and any remaining false positive cells were discarded. ABA RGB coordinates were then obtained from matching the new XY cell positions onto the corresponding transformed RGB atlas image obtained from the Atlas registration step, and the number of cells detected in a region were counted.

Axonal projection density was analyzed with custom written software (MATLAB Version R2018b, The MathWorks Inc.). Images were first de-noised using a Wiener-Filter (neighborhood size: 2x2). Image slice edges that displayed saturated signal due to histological processing were removed by edge correction from a corresponding binarized DAPI micrograph. Axons were then detected by convolving the images using the Maximum Response 8 (MR8) Gaussian filter bank (Varma and Zisserman, 2005) with a width of 3-6 pixels (corresponding to a minimum and maximum axon width of 1.2 µm and 2.4 µm respectively in the down sampled image). A single threshold for axon detection was determined empirically based on the length of the detect axon and was applied to all datasets. The same threshold value was then used for all the corresponding slices and associated datasets. Images were closely matched to the original micrographs to validate axon detection as well as identify and manually remove any residual noise pixels that appeared as a consequence of tissue processing (large, noncontiguous fluorescence). ABA RGB coordinates were then obtained by matching the XY pixel positions with the corresponding transformed RGB atlas image obtained during the Atlas registration step. Finally, we counted the number of pixels detected within a region.

#### Visualization

After atlas registration and signal detection, cell soma (input) and axons (output) were projected onto a 3D reference atlas in Imaris volumetric image software (Version 9.3, Bitplane AG), as previously described for the rat (Dempsey et al., 2017), using the matrix transformations for the ABA described in Puchades et al., 2019.

### Data analysis

Cells and axons were quantified across the whole brain of individual mice using custom-written Python code. Data were normalized as a fraction of the total amount of inputs or outputs detected across the brain. Data were grouped into 6 major brain regions (Cortex, Striatum/Pallidum, Amygdala, Thalamus/Hypothalamus, Midbrain, and Hindbrain) with 70 subregions as determined from the ABA.

S1 and pIC input-output Pearson correlation coefficients were performed for each major brain region (cortex, thalamus, and amygdala). Independent t-tests for each brain region were performed on the percentage of whole-brain inputs or outputs between S1 and pIC. All values are expressed as mean ± SEM unless otherwise stated. Statistically significant differences were considered at *p* < 0.05. No statistical methods were used to predetermine sample size.

To visualize the spatial alignment of the functional response and the injection location (Figure 1E, F – left), fluorescence contours were aligned across mice using custom-written Python code as described previously (Vestergaard et al., 2022). Briefly, the functional fluorescence images (cool- and sound-trial average evoked responses) and the anatomical fluorescence image (injection location) were smoothed with a Gaussian filter (σ = 20 pixels). The center-of-mass was computed from all pixels above a threshold of 80 % of peak fluorescence for the trial-averaged cool-evoked responses to identify the center of the cortical region sensitive to temperature. For visualization of the functional signal, 80^th^ percentile contours for each field of view (S1, pIC) from individual mice were translated to align to the center-of-mass of the fluorescence for thermal stimulation. For anatomical fluorescence images, the contours were superimposed on the corresponding atlas section (Figure 1E, F - right).

To assess spatial separability of the inputs or outputs within a given subdivision of the brain, the point cloud of coordinates for inputs (cell soma) and outputs (axon) within each subdivision were converted into a mesh in the ABA coordinate space. Contralateral coordinates were removed for this analysis. To minimize sampling limitations due to tissue thickness, the mesh was smoothed (spatial Gaussian, standard deviation = 100 µm). The mesh was converted to a binary matrix at a threshold value of one-tenth of the maximum voxel. A binary matrix was generated for S1 inputs, S1 outputs, IC inputs, and IC outputs for each brain subdivision. An overlap parameter was estimated for each subdivision as:

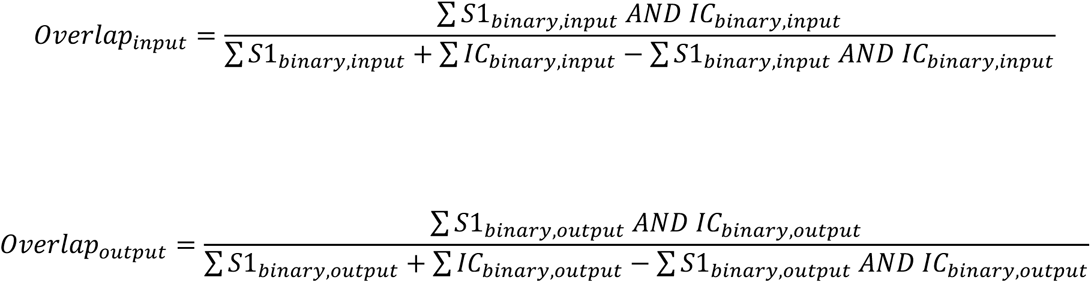

Random data sets (n = 50 per subdivision) were generated under the null hypothesis by shuffling the labeling of each coordinate included in the analysis for Monte Carlo hypothesis testing. The p-value was computed as the proportion of simulated overlap coefficients greater than the observed overlap coefficient.

To visualize the spatial overlap, the mesh was not binarized. Instead, a three-dimensional contour plot was generated (isosurface, Matlab) at the 30^th^ quantile of the non-zero voxels. As shown in Figure 6F and 6G, this spatial map was generated across three thalamic nuclei – VPL, POm, and POt.

**Figure 6.**
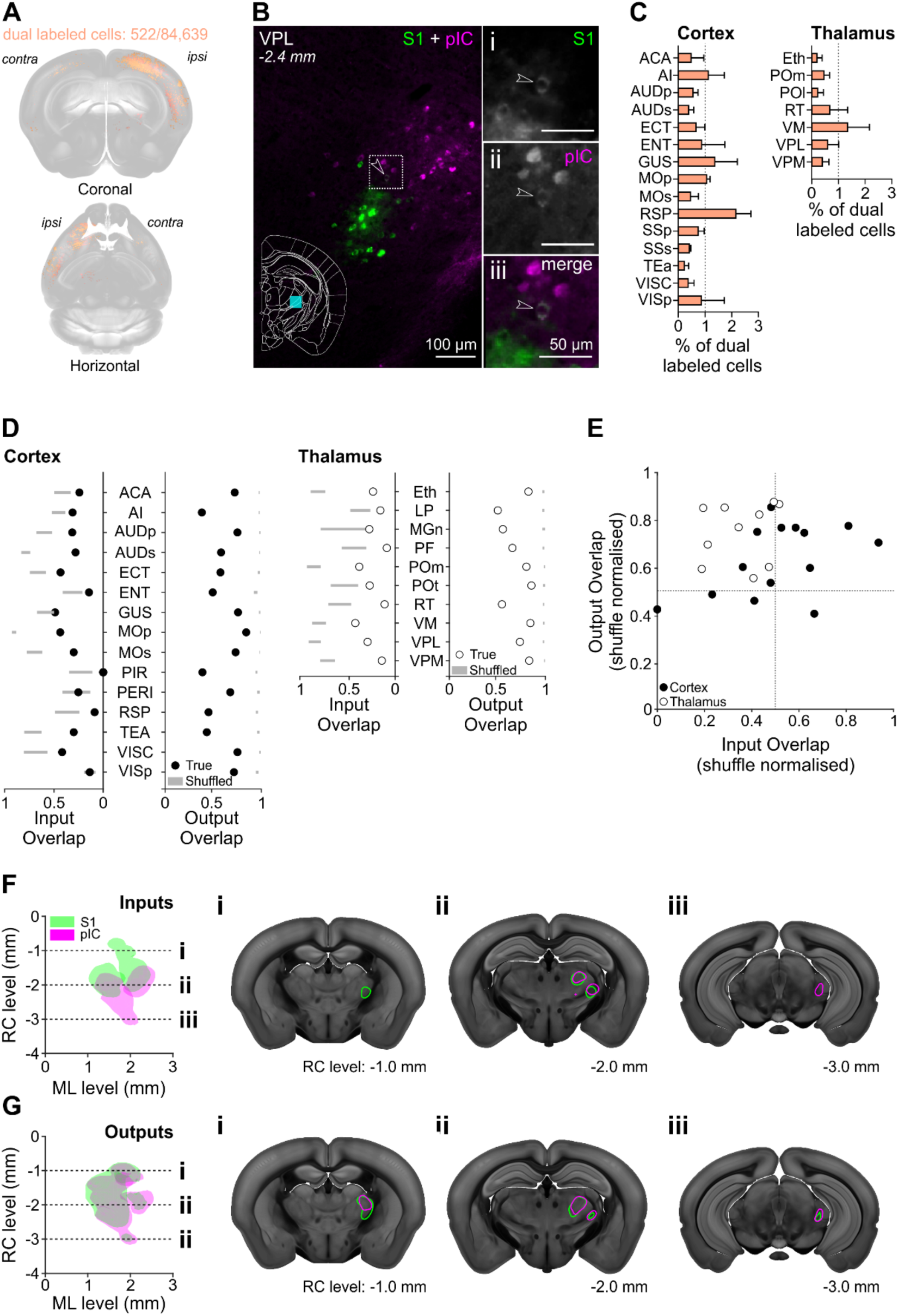
Spatial organization of cortical and thalamic inputs and outputs from S1 and pIC. **A**, Cell bodies labelled with both Ctb647 and Ctb555 (pseudocoloured orange) that project to S1 and pIC registered to the Allen CCF v 3.0. **B**, Representative micrograph of a coronal section showing the VPL nucleus with a Ctb positive cells projecting to S1 (**i** – green) or pIC (**ii** – magenta), and one identified cell that projects to both S1 and pIC (**iii** – white). **C**, Percentage of dual labelled cells projecting to S1 and pIC in cortical and thalamic nuclei (mean +/- SEM). Full list of abbreviations in Supplementary Table 1. **D**, Input and output overlap coefficients quantified from multiple cortical or thalamic nuclei. Dots show mean true data while gray bars represent 99 % confidence intervals of the shuffled distributions. **E**, Input and output overlap coefficients normalized by the shuffle distribution mean for each region. Each circle denotes an individual brain region. Open circles denote thalamic regions, close circles denote cortical regions. Detailed labelling of individual data points is provided in Supplementary Figure 8. The position of the data points in the upper left quadrant of the graph demonstrate that inputs are less overlapped than outputs. Note that thalamic regions (open circles) are further from the diagonal than cortical regions (black), indicating that the thalamic regions show a greater difference in their input/output spatial overlap than cortex. **F**, Horizontal projection of reconstructed VPL, POm, and POt inputs to S1 (green) or pIC (magenta) (left). Coronal sections showing a rostral targeting S1 region (**i**), intermediate S1 and pIC targeting region (**ii**) and a caudal targeting pIC region (**iii**). **G**, Same as F except for thalamic outputs from S1 (green) and pIC (magenta) showing major overlap at all rostral caudal axis (**i, ii, iii**).

### Data exclusion

Data were excluded if: (i) the injection site was not located in the cortical functional response; (ii) post-hoc examination showed that the injection site was mistargeted; (iii) if the retrograde injection spread into the underlying white matter tract (corpus callosum).

### Data and code availability

The datasets and code supporting the current study will be deposited to public repositories.

## RESULTS

### Functionally targeted tracer injections into forelimb S1 and pIC

We targeted the forelimb thermosensitive regions of S1 and pIC using wide-field calcium imaging in anesthetized GP 4.3 mice which express GCaMP6s in cortical excitatory neurons under the Thy1 promoter (Chen et al., 2013; Dana et al., 2014) (Figure 1B). The skull of the mouse was thinned and glass coverslips were mounted over the left-hand side of S1 and pIC. A 10 °C cooling stimulus (32 to 22 °C) was then delivered to the glabrous skin of the right forepaw via a Peltier element and evoked responses were visualized online (Figure 1C, D, left). The location of thermal representation in pIC was further confirmed by functional identification of the neighboring auditory cortex using an auditory tone (Figure 1D, F, left). Next, glass coverslips were removed and craniotomies drilled over the center of the thermal responses. Anatomical tracers were then injected into the center of the functional response via a glass micropipette (Figure 1B). Cholera-toxin b (Ctb) (555 or 647, pseudo colored magenta and green respectively) was used for retrograding labelling of cell bodies, and AAV2/1 (Ruby2sm-Flag or GFPsm-myc, pseudo-colored magenta and green respectively) for anterograde labelling of axonal outputs (Figure 1C, D). Finally, the spatial overlap between the functional response and injection was confirmed with in vivo imaging (Figure 1E, F, left).

Seven days after injection of Ctb, or three to four weeks after injection of AAVs, mice were perfused and brains removed for histological processing. Analysis of the Ctb and AAV injection sites in S1 and pIC showed that the spread of the tracers in the injection site was not significantly different and spread throughout the entire cortical column (medial/lateral pIC: 565 ± 40 µm, S1: 653 ± 77 µm; dorsal/ventral pIC: 927 ± 46 µm, S1: 986 ± 50 µm; rostral/caudal pIC: 800 ± 55 µm, S1: 800 ± 43 µm) (Figure 1E, F; Supplementary Figure 1). To identify brain wide input and output nuclei, fluorescent images of the whole brain were taken and registered to the Allen Brain Atlas Common Coordinate Framework v3 (ABA, Supplementary Figure 3). Regions 1 mm rostral/caudal and dorsal/ventral (parallel with the cortical region) from the center of the injection site were excluded from further analysis due to saturation of the fluorescent signal. DAPI staining of cell bodies and comparative analysis with the Scnn1a-tdTomato mouse line showed that the injections were targeted to a granular region of pIC (putative layer 4 thickness S1: 205 ± 14 µm and pIC: 215 ± 11 µm, Supplementary Figure 2).

Whole-brain input-output connectivity maps were created from coronal sections spaced 200 µm apart from +1.4 mm to -7.0 mm relative to bregma. Analysis of the olfactory bulbs, frontal cortical regions and the cerebellum were excluded. As the number of labelled neurons (S1: 5147 ± 800, pIC: 7737 ± 330 cells) and axons (S1: 2,323,883 ± 245,388, pIC:1,862,505 ± 286,668 pixels) varied across mice, the input and output values were normalized as a fraction of the total amount of input cell bodies/axonal outputs detected across the entire brain.

### Whole-brain input-output connectivity of forelimb S1 and pIC

To quantify the whole-brain inputs and outputs of S1 and pIC on a broad scale (Figure 2A, C), we initially defined 6 major regions; cortex, striatum/pallidum, amygdala, thalamus/hypothalamus, midbrain, and hindbrain (Figure 2B, D). The majority of inputs originated in the side ipsilateral to the injection site, with contralateral inputs almost exclusively located in the contralateral cortex (Figure 2A, S1 ipsi: 91 %, S1 contra: 9 %, pIC ipsi: 80 %, contra: 20 %). The overall brain-wide distribution of inputs was similar for S1 and pIC, with the cortex being the dominant source of ipsilateral inputs to S1 and pIC, however pIC showed a higher connectivity with the contralateral cortical regions than S1 (Figure 2B – right, S1: 8 %, pIC: 19 %, *p* = 0.01). The ipsilateral thalamus was the second major input region, with significantly more inputs projecting to S1 than pIC (Figure 2B - left, S1: 10 %, pIC: 6 %, *p* = 0.01). Intriguingly, the amygdala projected to pIC but not to S1.

**Figure 2.**
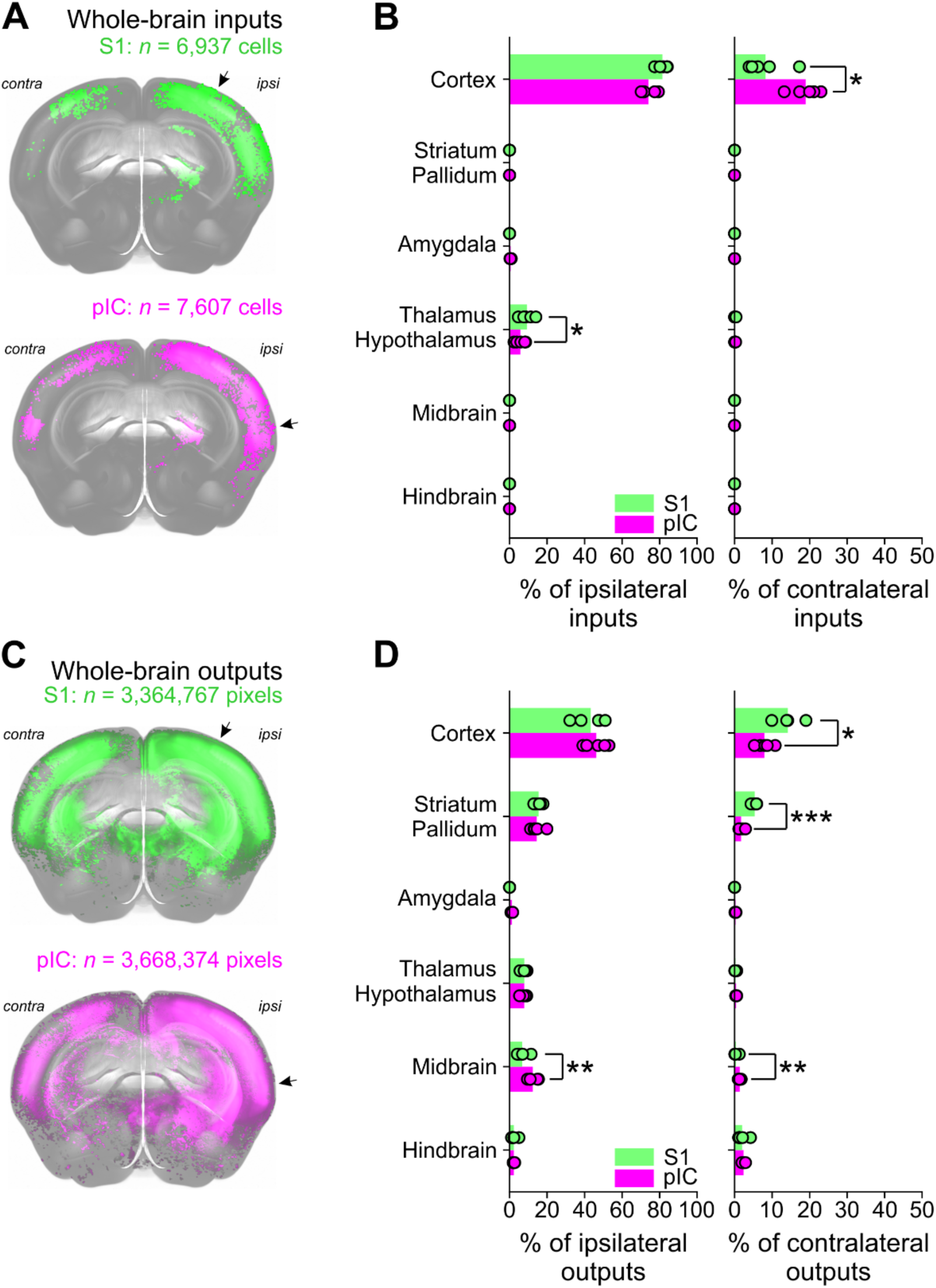
Whole-brain visualization of long-range inputs and outputs of thermal regions of S1 and pIC. **A**, Front view of a 3D brain reconstruction showing brain wide cell bodies providing input to S1 (top – green, n = 6,937 identified cell bodies) or pIC (bottom – magenta, n = 7,607 identified cell bodies), from one example mouse. Arrows indicate the location of injection. **B**, A comparison of the ipsilateral (left) and contralateral (right) inputs from 6 brain regions to S1 (green) or pIC (magenta) as a percentage of the whole-brain inputs. * shows significant difference between inputs to S1 vs pIC (n = 5 mice, p < 0.05, two-way independent *t*-test). Bars show means, open circles show individual mice. **C**, Same as **B** but showing reconstruction of pixels labelled with axonal outputs (green, n = 3,364,767 pixels; magenta n = 3,668,373 pixels). **D**, Same as **B**, but showing a comparison of cortical axonal outputs in target regions (* = p < 0.05, ** = p < 0.01, *** = p < 0.001).

Like inputs to S1 and pIC, brain-wide outputs mostly targeted regions ipsilateral to the injection site (Figure 2B, S1 ipsi: 74 %, S1 contra: 26, pIC ipsi: 86 %, pIC contra: 14 %). The major target of S1 and pIC axons was the cortex, which received a similar amount of ipsilateral innervation (Figure 2D - right, S1: 43 %, pIC: 46 %, *p* = 0.11), while projections from S1 to the contralateral cortex were stronger than from pIC (Figure 2D – left, S1: 14 %, pIC: 8 %, *p* = 0.01). The second major innervation target of S1 and pIC was the striatum/pallidum, which received similar levels of ipsilateral input (Figure 2D - left, S1: 15 %, pIC 14 %, *p* = 0.6), but significantly more contralateral input from S1 than pIC (Figure 2D - right, S1: 5 %, pIC: 2 %, *p* = 0.001). Ipsilateral and contralateral axonal targets from S1 and pIC innervated the thalamus (Figure 2D, S1 ipsi: 8 %, pIC ipsi: 8 %, *p* = 0.98, S1 contra: 0.2 %, pIC contra: 0.4 %, *p* = 0.19) and hindbrain (Figure 2D, S1 ipsi: 2 %, pIC ipsi: 2 %, *p* = 0.98, S1 contra: 2 %, pIC contra: 2 %, *p* = 0.52) to similar levels. Both ipsilateral and contralateral sides of the midbrain received more innervation from pIC compared to S1 (S1 ipsi: 7 %, pIC ipsi: 12 %, *p* = 0.004, S1 contra: 0.3 %, pIC contra: 1 %, *p* = 0.002). Closely resembling the inputs, the amygdala was innervated exclusively by pIC outputs and not by S1 (pIC ipsi: 1.2 %, pIC contra: 0.2 %).

To examine connectivity at higher resolution, we went on to subdivide the 6 major anatomical areas into 70 subregions and present data from regions ipsilateral to the injection side (for a comprehensive list of all ipsilateral and contralateral connections, see Supplementary Figures 4, 5, and Supplementary Tables 2, 3). To assess S1 and pIC connectivity at different scales, we compare the connectivity strength as well as spatial distributions of regions, subregion nuclei and single cells.

**Figure 4.**
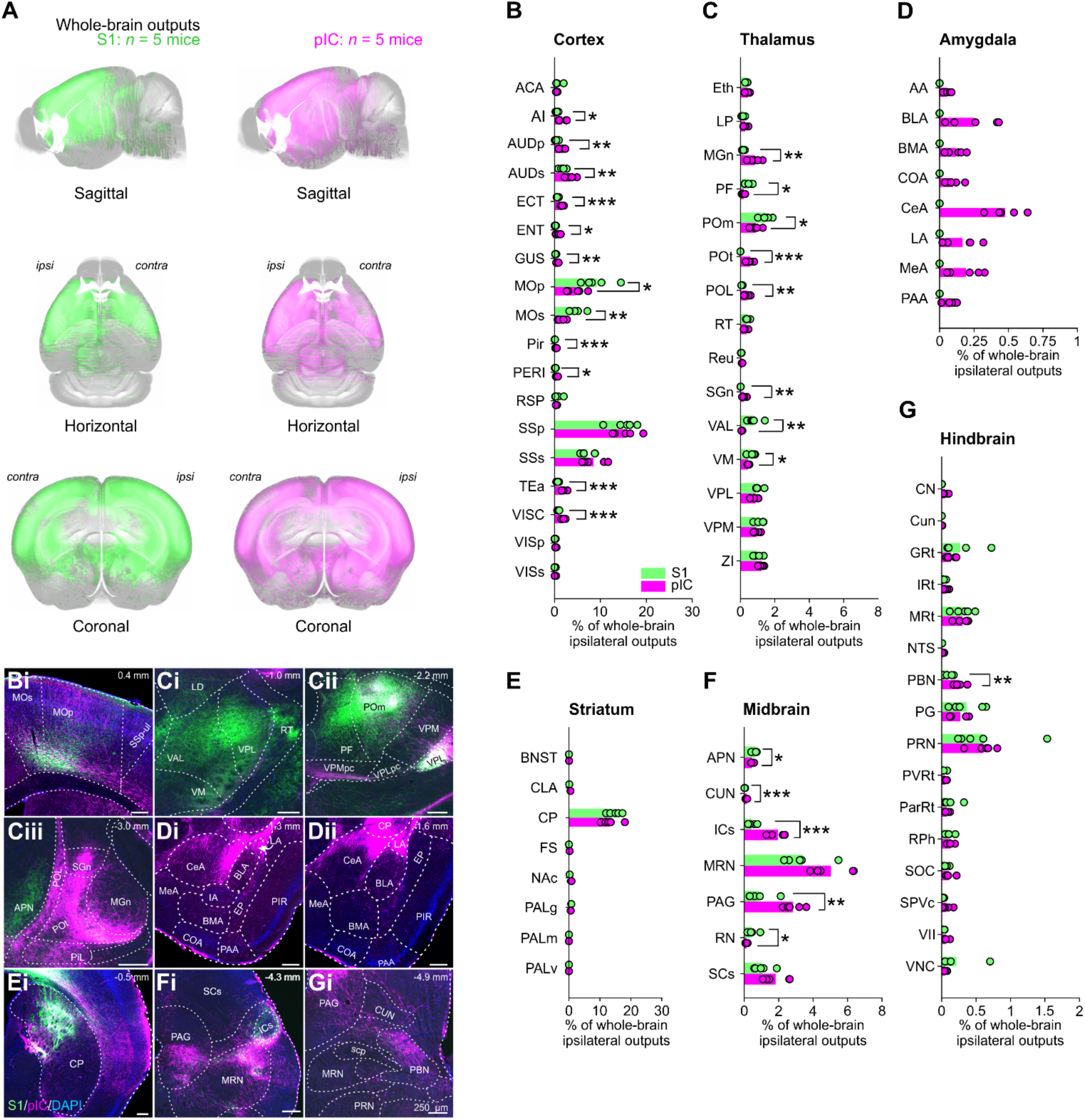
Whole-brain ipsilateral outputs from thermally responsive S1 and pIC. **A**, AAV fluorescence viruses were injected into thermally responsive areas of S1 and pIC. The identified axons projecting from S1 (left) or pIC (right) were extracted and registered to the Allen CCF v 3.0 **B, C, D, E, F, G**, Proportions of whole-brain ipsilateral outputs from S1 (green), or pIC (magenta) across individual (**B**) cortical, (**C**) thalamic, (**D**) amygdaloid, (**E**) striatal, (**F**) midbrain, or (**G**) hindbrain subregions. Bars show means and open circles show individual mice. * = p < 0.05, ** = p < 0.01, n = 5 mice per condition. See Supplementary Table 2 for proportions of all subregions. Representative example brain slices of outputs from S1 (green) or pIC (magenta) from different (**Bi**) cortical, (**Ci, Cii, Ciii**) thalamic, (**Di, Dii**) amygdaloid, (**Ei**) striatal, (**Fi**), midbrain (**Gi**) and hindbrain subregions. Scale bars, 250 µm. A list of abbreviations is provided in Supplementary Table 1.

**Figure 5.**
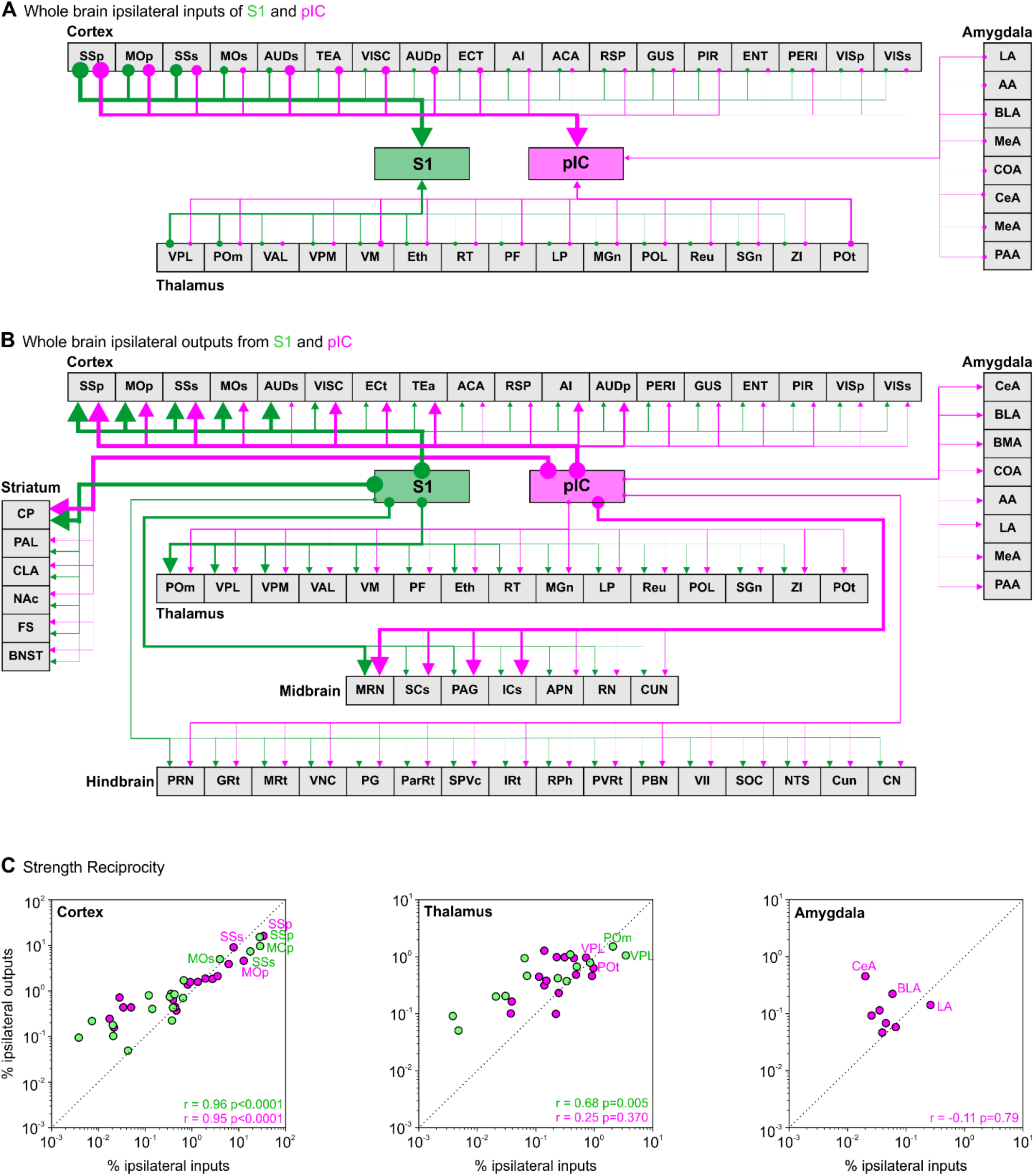
Wiring diagrams and reciprocity of the strength of inputs and outputs to S1 and pIC. **A**, Schematic wiring diagram showing regions that project to S1 (green) or pIC (magenta). Regions are ordered from left to right based on their projections to S1. Line thickness corresponds to the strengths of inputs (low, medium, high) in Figure 3 and Supplementary Table 2. For a list of subregion abbreviations see Supplementary Table 1. **B**, Same as (A) but for axonal projections. Line thickness corresponds to the amount of outputs (low, medium, high) observed and indicated in Figure 4 and Supplementary Table 3. **C**, Correlations of input/output strength for S1 (green) and pIC (magenta) subregions in (top) cortex, (middle) thalamus, and (bottom) amygdala. Individual data points correspond to subregion mean value across mice (r = Pearson’s correlation coefficient). Detailed input/output strength correlations including labelling of individual data points is provided in Supplementary Figure 7.

### Whole-brain inputs to forelimb S1 and pIC

Visualizing inputs to S1 and pIC at different angles of a 3D projection, revealed dense labelling across many cortical and thalamic nuclei (Figure 3A, Supplementary Table 2, Supplementary Movie 1). The majority of inputs showed a similar innervation strength to S1 and pIC. Notable exceptions included stronger inputs to S1 than pIC from regions involved in sensorimotor processing, including primary motor cortex (MOp, 26 %, *p* = 0.03, Figure 3B, 3Bi) and secondary somatosensory cortex (SSs, 21 %, *p* = 0.01). In support of its role in diverse sensory and cognitive functions (Gogolla, 2017), pIC received significantly input from a broader range of cortical nuclei, including agranular insular cortex (AI, 0.8 %, *p* = 0.02), primary and dorsal/ventral auditory cortices (AUDp, 2.5 %, *p* = 0.002, AUDs: 5.4 %, *p* = 0.009), retrosplenial cortex (RSP, 0.4 %, *p* = 0.03), temporal association area (TEA, 3.4 %, *p* = 0.04), and visceral cortex (VISC, 2.2 %, *p* = 0.004). S1 received significantly more thalamic input from nuclei within the ventral basal and posterior thalamic compartments (Figure 3C, Ci, Cii), including the ventral posterolateral (VPL and its parvocellular compartment VPLpc, 4 %, *p* = 0.01), the ventral anterolateral (VAL, 0.7 %, *p* = 0.04), and posterior medial (POm, 2.6 %, *p* = 0.02, Figure 3Cii). Thalamic innervation of the pIC was more diverse than S1, with significantly more input arising from the medial geniculate nucleus (MGn, 0.5 %, *p* = 0.002) and a prominent innervation from the primary triangular (POt) nucleus (Figure 3Ciii, 1.1 %, *p* = 0.006) that did not project to S1. In agreement with prior literature (Shi and Cassell, 1998a, 1998b; Schiff et al., 2017), we did not observe any innervation of S1 by the amygdala (Figure 3D). In contrast, pIC was innervated by cortical-like regions of the amygdala, the majority of which came from the lateral amygdala (LA, 0.3 %, Figure 3Di). The baso-lateral (BLA, 0.06 %, Figure 3Dii) and piriform-amygdala area (PAA, 0.07 %, Figure 3Dii) innervated pIC to a lesser extent and very few retrogradely labelled cell bodies from S1 and pIC were observed in the striatum-like centromedial nuclei (CeA, MeA both < 0.05 %). Together these data show that pIC and S1 share input regions with the exception of the amygdala that projects exclusively to pIC.

**Figure 3.**
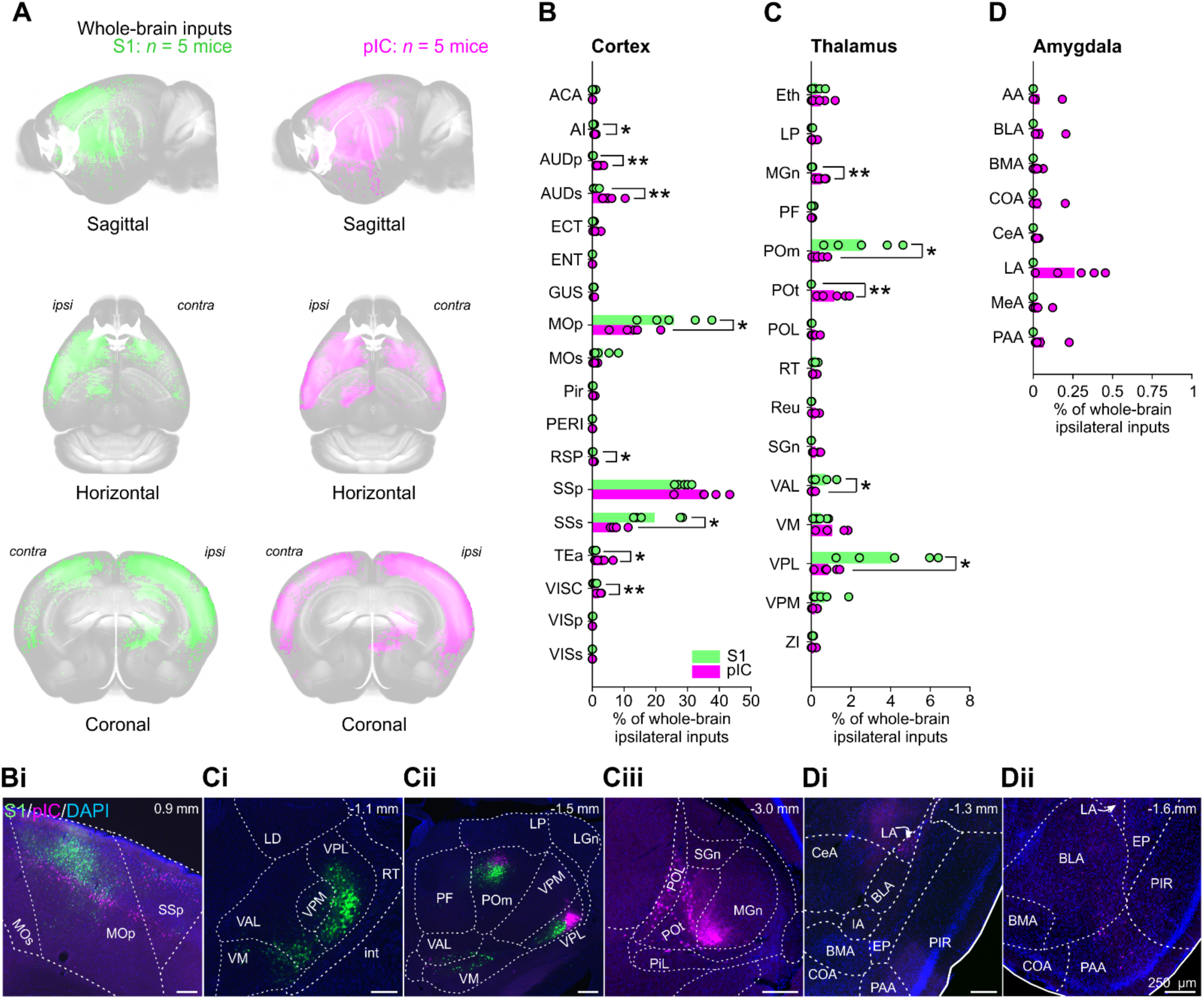
Whole-brain ipsilateral inputs to thermally responsive in S1 and pIC. **A**, Whole brain 3D images of cell bodies labelled with CTb 647 (pseudo-coloured green, S1 injection) or Ctb 555 (pseudo-coloured magenta, pIC injection). Cell bodies were identified and registered to the Allen CCF v 3.0. **B, C, D**, Proportions of whole-brain ipsilateral inputs to S1 (green), or pIC (magenta) (**B**) cortical, (**C**) thalamic, or (**D**) amygdaloid subregions. Bars show means and open circles show individual mice. * = p < 0.05, ** = p < 0.01, n = 5 mice per condition. See Supplementary Table 2 for proportions of all subregions. Representative example brain slices of inputs to S1 (green) or pIC (magenta) from different (**Bi**) cortical, (**Ci, Cii, Ciii**) thalamic or (**Di, Dii**) amygdaloid subregions. Scale bars, 250 µm. A list of abbreviations is shown in Supplementary Table 1.

### Axonal output targets of forelimb S1 and pIC

Axonal projections from S1 and pIC broadly innervate the entire mouse brain (Figure 4A), with S1 and pIC both having more output than input targets. At subregion resolution, we noted a number of significant differences in the comparative strengths of innervation (Figure 4B to F, Supplementary Table 3, Supplementary Movie 2). Similar to the pattern of inputs to S1 and pIC, axonal outputs from S1 mostly innervated regions involved in somatosensory processing, such as primary and secondary motor cortices (MOp, 9 %, *p* = 0.03, MOs, 5 %, *p* = 0.006, Figure 4Bi), whereas pIC neurons innervated a broader number of cortical regions areas, including AI (1.6 %, *p* = 0.02), AUDp (1.6 %, *p* = 0.005), AUDs (3.5 %, *p* = 0.01), ectorhinal (ECT, 1.6 %, *p* = 0.0004), entorhinal (ENT, 0.8 %, *p* = 0.02), gustatory (GUS: 0.7 %, *p* = 0.004), piriform (Pir, 0.4 %, *p* = 0.0006), perirhinal (PERI, 0.5 %, *p* = 0.02), TEA (2 %, *p* = 0.0009) and VISC (1.9 %, *p* = 0.0005). Notable differences between the thalamic innervation from S1 and pIC included significantly more S1 outputs innervating VAL (0.7 %, *p* = 0.003, Figure 4Ci), the ventral medial nucleus (VM, 0.7 %, *p* = 0.04, Figure 4Ci), the parafascicular nucleus (PF, 0.5 %, *p* = 0.02, Figure 4Cii), and POm (1.5 %, *p* = 0.01, Figure 4Cii); while stronger pIC innervation was observed in MGn (0.8 %, *p* = 0.004), the suprageniculate nucleus (SGn, 0.02 %, *p* = 0.003), and the posterior limiting nucleus (POL, 0.4 %, *p* = 0.002). As with their inputs, the POt was selectively innervated by pIC and not by S1 (0.6 %, Figure 4Ciii). Similarly, the amygdala was innervated by pIC and not by S1 (Figure 4D) with the majority of pIC outputs targeting the striatum like central and medial amygdala nuclei (CeA and MeA, collectively 0.67 %, Figure 4Di, 4Dii), and to a lesser extent, the cortical like basolateral and basomedial nuclei (BLA, BMA and LA, collectively 0.54 %, Figure 4Di, 4Dii). Unlike their inputs, outputs from S1 and pIC strongly targeted a range of nuclei in the midbrain and hindbrain. The primary target of both S1 and pIC was the MRN, however S1 axons significantly innervated the anterior pretectal nucleus (APN, 0.45 %, *p* = 0.04, Figure 4Ciii) and the red nucleus (RN, 0.13 %, *p* = 0.04) more than pIC (Figure 4E), whereas pIC showed significantly more innervation of the cuneiform nucleus (CUN, 0.13 %, p = 0.0006), inferior colliculus (ICs: 2 %, *p* = 0.0002, Figure 4Ei), and the periaqueductal gray (PAG, 2.8 %, *p* = 0.001, Figure 4Ei). The hindbrain received similar levels of axonal output from both S1 and pIC (Figure 4F), with one notable difference being the stronger innervation of the parabrachial nucleus by the pIC (PBN, 0.25 %, *p* = 0.006, Figure 4Fi), an area that forwards thermal information to circuits in the hypothalamus that regulate body temperature.

### Reciprocal connectivity of forelimb S1 and pIC

A canonical feature of cortical wiring is reciprocal connectivity, whereby regions providing input also receive outputs from the target region. Plotting the input and output circuit diagrams from our tracing data highlighted that a number of cortical and thalamic regions were reciprocally connected with S1 and pIC (Figure 5A, B). In contrast, the hindbrain, midbrain and striatum only received axonal projections from S1 and pIC without providing direct inputs. To investigate the reciprocal connectivity between S1 and pIC, we plotted the strengths of cortical, thalamic and amygdaloid inputs against their respective innervation from S1 and pIC (Figure 5C, Supplementary Figure 7). In agreement with an established model of cortico-cortical connectivity (Felleman and Van Essen, 1991), we observed strong reciprocity for both S1 and pIC with other cortical regions (S1: r = 0.96, *p* < 0.0001, pIC: r = 0.95, *p* < 0.0001). The thalamus was highly reciprocal with S1 ((r = 0.68, *p* = 0.005) and not with pIC (r = 0.25, *p* = 0.370). The connectivity between pIC and the amygdala was dominated by outputs from the pIC, and, while it did provide inputs to pIC, the correlation between input and output strength was not significant at the population level (r = -0.11, *p* = 0.79).

### Spatial organization of S1 and pIC whole-brain inputs and outputs

On a subregion level, S1 and pIC have similar input structures and output targets. However, this does not address whether there was target specific connectivity of individual cells (parallel pathways) or an absence of distinct substructure organization (mixed pathways). To assess whether subregions contained cells projecting to both S1 and pIC, cells labeled with both retrograde tracers were counted and projected onto the 3D mouse brain. Dual labelled neurons were sparse (Figure 6A) compared to the total inputs innervating S1 or pIC. Of all the 84,639 total cells (5 mice) that projected to S1 or pIC, we identified only 522 neurons that were dual labelled (Figure 6A, 6Bi, Bii). From subregions providing input to both S1 and pIC (Figure 6C) only 0.65 % cortical, and 0.36 % thalamic neurons projected to both, together indicating that input is provided by separate circuits.

As the inputs to S1 and pIC arose primarily from separate populations, we went on to analyze the spatial organization of clusters of inputs within each cortical and thalamic subregion. We defined spatial separability, or overlap (see methods), as the percent of voxels containing both S1 and pIC projecting neurons within a given subregion. A completely separable map like organization would have an overlap value of 0, while a completely intermingled salt-and-pepper like organization would have an overlap value of 1. We found that within the cortex and thalamus, most co-labelled subregions showed non-random, spatially organized inputs that were significantly different from their shuffled distributions (Figure 6D). This indicates that inputs from the cortex and thalamus to S1 and pIC are organized in a spatially separate map-like arrangement within individual subregions.

We next asked whether this was also true for axonal projections from S1 and pIC. We found that cortical and thalamic regions had less spatial overlap of S1 and pIC axons than expected from a random distribution (Figure 6D). However, while both inputs and outputs demonstrated spatial separability, the separation for inputs is much higher than outputs (average input overlap: 0.25, average output overlap: 0.67, Figure 6D). Even after normalizing the overlap value to the shuffled control value, the input overlap is smaller than the output overlap across cortical and thalamic subregions (Fig 6E, Supplementary Figure 8). This can be visualized most clearly in thalamic regions where there is a clear rostrocaudal division in both the input and output volumes (Fig 6F). In a horizontal projection of VPL, POm, and POt, the S1 inputs are spatially localized to the rostral region of thalamus, while the pIC inputs are spatially localized to the caudal regions. At more rostral levels, coronal slices contain exclusively S1 projecting cells (Figure 6Fi) while caudal coronal slices contain exclusively pIC projecting cells (Figure 6Fiii) with some overlap at intermediate rostrocaudal levels (Figure 6Fii). In the corresponding output plots, there is a significantly different distribution of S1 and pIC axonal outputs but a reduced separability compared to thalamic inputs (Figure 6G).

While S1 and pIC share multiple input and output nuclei at a gross scale, these results suggest that there are finer patterns of connectivity within cortical and thalamic subregions. The input neurons projecting to S1 and pIC arise from spatially separate populations within each subregion, supporting the hypothesis that S1 and pIC have parallel input pathways. Though the output projections of S1 and pIC showed non-random spatial separability within each subregion, this separation was lower than the input populations, supporting a more mixed model of S1 and pIC outputs.

## DISCUSSION

Here we used functionally targeted tracer injections to generate a comprehensive map of long-range inputs and outputs from two cortical representations of temperature. This approach allowed a direct comparison of S1 and pIC connectivity in the same mice. While both areas receive input from common cortical and thalamic regions, pIC thalamic input is more widespread and, at cellular resolution, inputs to both areas originated from largely non-overlapping and spatially separated neuronal populations. Despite receiving independent inputs, S1 and pIC innervate similar long range cortical and subcortical regions with axonal projections that were less spatially separated than their inputs, implying that the formation of coherent thermal percepts involves the convergence of cortical outputs. Notably, exclusive connectivity was observed between the pIC and the amygdala, PBN of the hindbrain, and the POt. Together, our data suggest that thermal information forms at least two separate pathways that run via S1 and pIC and are widely broadcast across the brain.

### Limitations and considerations – identification and nomenclature of pIC

A classic approach to address wiring of a brain region is to inject neuronal tracers using bregma coordinates for targeting a region of interest. Bregma coordinates, however, are notoriously variable from mouse to mouse. To address this, we functionally targeted our injections to the center of the widefield cortical calcium response to cool stimuli delivered to the forepaw. In order to standardize connectivity maps between mice, we then aligned brain slices to the mouse brain atlas from the Allen Mouse Brain Common Coordinate Framework version 3, which is widely used to standardize maps of neural circuitry (ABA, Pollak Dorocic et al., 2014; Do et al., 2016; Niedworok et al., 2016; Zhang et al., 2016; Fürth et al., 2018; Wang et al., 2020; Benavidez et al., 2021; Dempsey et al., 2021). Aligning to the ABA allowed correction for multi-plane distortion, misalignment, and tissue deformation during tissue processing, however, area borders in the ABA have chiefly been constructed using anatomical markers rather than functional properties of regions, potentially leading to discrepancies in less well studied areas.

In this study, functionally targeted injections into the thermal region of forelimb S1 were localized post-hoc in the primary somatosensory upper limb region of the ABA (SSp-ul). Injections into pIC, however, were labelled as secondary somatosensory cortex (SSs) or the dorsal and ventral auditory area (AUDs, see: Figure 1). Work in multiple mammalian species has identified a somatotopic representation of cutaneous tactile input located in an area ventral to secondary somatosensory cortex (SSs), dorsal to the rhinal vein, bordering rostral regions of the auditory cortex and containing an anterior auditory field and termed parietal ventral (PV) area or posterior insular cortex (pIC) (in squirrels: Krubitzer et al., 1986, hedgehog: Catania et al., 2000, opossum: Beck et al., 1996, marmosets/macaques: Krubitzer and Kaas, 1990; Krubitzer et al., 1995, mice: Gogolla et al., 2014; Nishimura et al., 2015, rat: Fabri and Burton, 1991a; Remple et al., 2003; Rodgers et al., 2008; Zhang et al., 2020). Using widefield calcium imaging, we have recently shown that this area contains somatotopically organized and rich cellular representations of cool and warm (Vestergaard et al., 2022), in contrast to the cool dominated representation in S1. Together with its profound impact on thermal perception, these data support the hypotheses that this region houses the primary cortical representation of temperature.

Recently, Gămănuţ et al. (2018) generated a new horizontal cortical map using multiple histological staining methods including transgenic mice expressing fluorophores (tdTomato) in parvalbumin neurons and muscarinic acetylcholine receptor, as well as cytochrome oxidase and VGlut2 staining. In this map, the borders of the visceral cortex (VISC), a cortical region likely homologous to granular IC, are significantly different to that of the ABA (see Supplementary Figures 1 and 3 of Gămănuţ et al., 2018), with VISC forming a lip that envelopes SSs and extends to the dorsal auditory area (AUDd) rather than ending abruptly and giving rise to the temporal association area (TEa) and ventral auditory area (AUDv) as in the ABA. In our recent work (see Supplementary Figure 1 of Vestergaard et al., 2022), alignment of flattened cortical sections to the horizontal cortical atlas modified from Gămănuţ et al., 2018 showed that the temperature response region in pIC was localized in VISC and agrees well with their separation of the S2 boundary with VISC. Together these data suggest that a granular portion of IC (or ‘VISC’) runs rostro-caudally along the dorsal aspect of anterior IC. Comparing our injection site locations to the same coronal sections from the Scnn1a mouse line, which labels layer 4 cortical neurons (Madisen et al., 2010), suggested that pIC was localized within a granular region of IC (Supplementary Figure 2).

While prior work has named this area PV or pIC, given their tightly overlapping location and similar sensory representation, we support the proposal by Rodgers et al., 2008 and Nishimura et al., 2015 that PV and pIC are in fact homologous areas. In agreement with the naming used by prior work in rodents (Rodgers et al., 2008, 2008; Gogolla et al., 2014; Beukema et al., 2018; Zhang et al., 2020) and evidence of thermal information processing in human posterior insular cortex (Craig et al., 2000), here we use the term pIC. To resolve this issue further and confirm the borders between AUDv, VISC, SSs (as according to the ABA), future work should perform detailed functional somatotopic mapping of thermal and tactile responses in pIC and SSs using acoustic stimuli to mark the insular auditory field and AUDv and AUDp followed by post-hoc histological mapping.

### Limitations and considerations – neuronal tracing methods

A critical limiting factor in the interpretation of tracing data lies in the spatial and functional specificity of the labeling methods. For example, tracer injections could still have spread into neighboring cortical regions like the primary motor cortex or the somatosensory barrel field that borders forelimb S1. We addressed this by measuring the spread of tracer for each experiment and noted that the measured spread of the injectant was unlikely to have spread into neighboring regions. However, the functional diversity within S1 could not be avoided. The core motivation of this study was to map and compare central circuits involved in thermal processing, but tactile and thermally responsive cells are spatially intermingled in S1, and are closely neighboring in pIC (Milenkovic et al., 2014; Vestergaard et al., 2022). Furthermore, pIC contains a thermal representation and a small auditory representation, the insular auditory field (Sawatari et al., 2011; Vestergaard et al., 2022). Therefore, our CTb and AAV based tracing from thermally responsive cortical regions would have labeled touch, temperature and, in pIC, possibly auditory responsive cells. In future work, this could be addressed by labelling neurons activated only by specific sensory stimuli through activity dependent expression of fluorescent proteins (e.g. Beukema et al., 2018) or methods like CANE or TRAP (Capturing and manipulating Activate Neural Ensembles, and Targeted Recombination in Active Populations) that can be coupled with anterograde or retrograde tracers (Guenthner et al., 2013; Sakurai et al., 2016; Wall et al., 2019). Single cell electroporation of functionally identified cells could also help investigate thermosensory specific neuronal circuits, but would be limited to very few cells (Wickersham et al., 2007; Marshel et al., 2010; Rancz et al., 2011). Despite these limitations, injections shown here encompass both S1 and pIC thermal representations with the data providing a roadmap for future functional and anatomical studies.

Our study also shared limitations common to anatomical tracing studies (Saleeba et al., 2019). First, the dense expression and bright fluorescence signal from both CTb and AAV labelling ∼1 mm^3^ from the center of the injection site prevents the analysis of local connectivity and we therefore masked this region from analysis. Second, though the direction of transport of CTb is primarily retrograde, it can also label in an anterograde direction. In our datasets, and only at high illumination for signals close to saturation, we observed some anterograde CTb+ve axons innervating the striatum (data not shown), but as our automated input analysis was tuned to identify cell soma (see Methods), these axons were left undetected and discarded upon manual confirmation. Third, CTb can be taken up by fibers of passage rather than terminating axons or cell bodies (Chen and Aston-Jones, 1995). The use of trans-synaptic retrograde tracing strategies, including rabies virus based tracing (Wickersham et al., 2007), retrograde AAVs (Tervo et al., 2016), or herpes simplex virus 1 (HSV-1, Ugolini et al., 1987), may help address this issue in future studies. However, these methods have their own caveats including tropism for cortical specific layers and the cellular mechanisms of trans-synaptic transport remain unclear. Moreover, due to the high connectivity seen between S1 and pIC here, dual rabies based technologies could label both populations equally and therefore starter populations would not be localized to either S1 or pIC. Lastly, while AAV based labelling is highly effective in labelling axonal projections, it does not specifically label synaptic boutons and therefore analysis of axonal projections will include fibers of passage as well as terminating fibers. Future use of tools that also label presynaptic sites, e.g. synaptophysin-cre based viruses (Beier et al., 2015; Lerner et al., 2015; Knowland et al., 2017; Dempsey et al., 2021) could help to resolve this.

### Comparison to previous anatomical tracing studies of forelimb S1 and pIC

Despite these possible limitations, our data show similar overall connectivity to previous connectivity mapping attempts from rodent forepaw S1 (Fabri and Burton, 1991a, 1991b, Zingg et al., 2014). For example, we observed that inputs to S1 originated from cortical somatosensory (SSp, SSs) and motor regions (MOp) as well as lemniscal and paralemniscal thalamic nuclei (VPL, POm). Moreover, S1 targeted similar cortical, striatal, thalamic, mid- and hind-brain subregions to the whole-brain forepaw S1 output mapping by Zakiewicz et al., 2014. Two differences to Zakiewicz et al., 2014 were strong projections from S1 to PAG and MRN and the lack of innervation of the substantia nigra. These differences could result from the use of different model systems (rat vs mouse), different tracers (BDA vs AAV), or injection volumes and spread. In support of this, a similar approach using virally expressed tracers by Oh et al., 2014 showed axonal projections from forelimb S1 to PAG and MRN, whereas only faint projections were observed following anterograde tracer injections of Phaseolus vulgarisleukoagglutinin (Pha-L) or biotinylated dextran amine (BDA) in Zingg et al., 2014.

The connectivity of rodent insular cortex has received great attention (Akers and Killackey, 1978; Guldin and Markowitsch, 1983; Cechetto and Saper, 1987; Allen et al., 1991; Shi and Cassell, 1998a, 1998b; Kimura et al., 2010; Mathiasen et al., 2015; Gehrlach et al., 2020), but in the majority of these studies tracer injections were not functionally targeted and only a small minority partially labelled the pIC region we examine here. Detailed comparisons are therefore challenging, but in broad agreement with prior work, we found that somatosensory and associated motor cortices (SSp, SSs and MOp) as well as key somatosensory thalamic nuclei (VPL, POm, POt) and amygdaloid subregions (LA, PAA and BLA) provide input to pIC. Likewise output targets were similar to those reported by (Shi and Cassell, 1998a) and (Kimura et al., 2010) who targeted the rat insular auditory field using electrophysiological mapping. One difference to Kimura et al. 2010, was the projection from pIC to the midbrain PAG, however in agreement with our data lateral PAG innervation has been observed from a caudal granular insular cortex region in mice (Oh et al., 2014; Zingg et al., 2014), perhaps reflecting a technical or species difference.

### Outlook

Understanding the wiring of the thermal system is important for a mechanistic assessment of thermal perception as well as the binding of diverse thermo-tactile percepts. Our results indicate that thermal information is routed to S1 and pIC by non-overlapping input pathways. It remains unclear why there is a far weaker representation of warm in forelimb S1 compared to pIC (Vestergaard et al., 2022), however, the identification of separate input pathways could help address this unexpected functional difference. From forelimb S1 and pIC, sensory information is then forwarded to more overlapping areas, perhaps reflecting the need to bind complex features during haptic exploration.

We observed exclusive connectivity of pIC with the amygdala, parabrachial nucleus (PBN), and POt. The PBN is known to forward thermal information to the hypothalamus for the control of body temperature (Madden and Morrison, 2019) and this connection suggests that pIC could provide top-down influence on body temperature. The POt receives direct input from dorsal layers of the spinal cord (Gauriau and Bernard, 2004; Al-Khater et al., 2008), and has been proposed to be the rodent homologue of VMpo (Gauriau, 2004), a structure in the primate thalamus proposed to be responsible for the routing of thermal information to pIC (Craig et al., 1994). Forelimb S1 and pIC also share connectivity with other thalamic nuclei, and a key question for future work is to address the cellular encoding of non-painful temperature in the thalamus. The strong connectivity of pIC with amygdala supports a role for valence encoding of temperature, and may help address the mechanisms behind the rapid learning of thermal perception tasks in mice (Paricio-Montesinos et al., 2020). Overall, our study provides a framework for the identification of neural mechanisms of thermal perception.

## Acknowledgements

We thank Alison Barth, Niccolo Zampieri, and members of the Poulet lab for constructive comments on earlier versions of the manuscript, and Svenja Steinfelder for help with administrative and technical aspects.

## Author contributions

P.B., and J.F.A.P designed the study. P.B performed all experiments. P.B., C.J.W., and T.M.L analyzed the data. P.B. and J.F.A.P wrote the manuscript.

## Data and material availability

Datasets and code will be deposited on public repositories.

