## Supplementary Information for "Brain-wide connectivity map of mouse thermosensory cortices"

**Supplementary material:** 22 pages including 7 Figures, 3 Tables, 2 Movies, and Supplementary References.

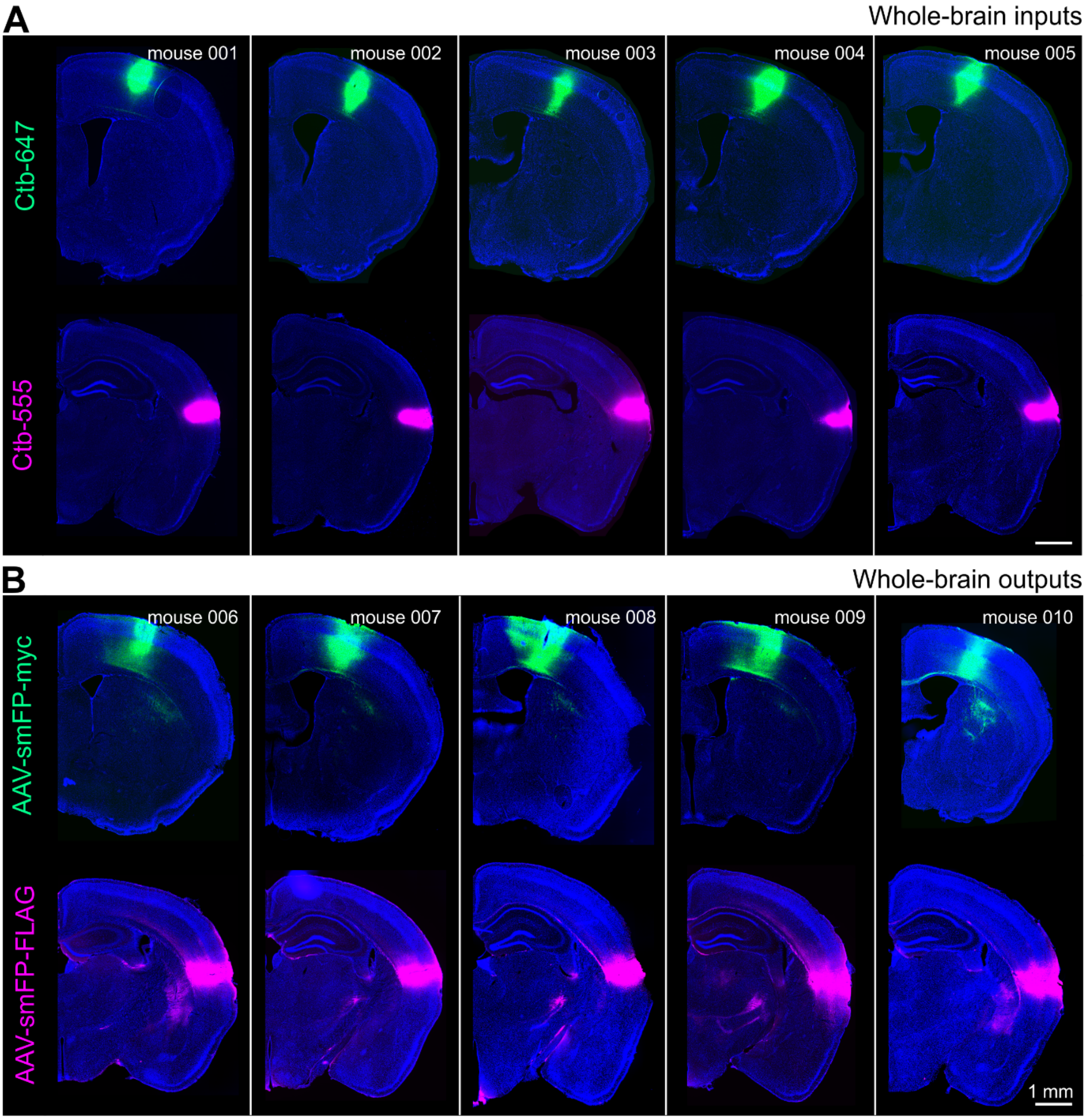


**Supplementary Figure 1. Example brain slices showing injection sites from all mice in study.**

**A**, Injection sites of CTb into thermal representation in S1 (green, top) or pIC (magenta, bottom) for input mapping. **B**, As above except for injection of anterograde AAV tracer sites for output mapping.

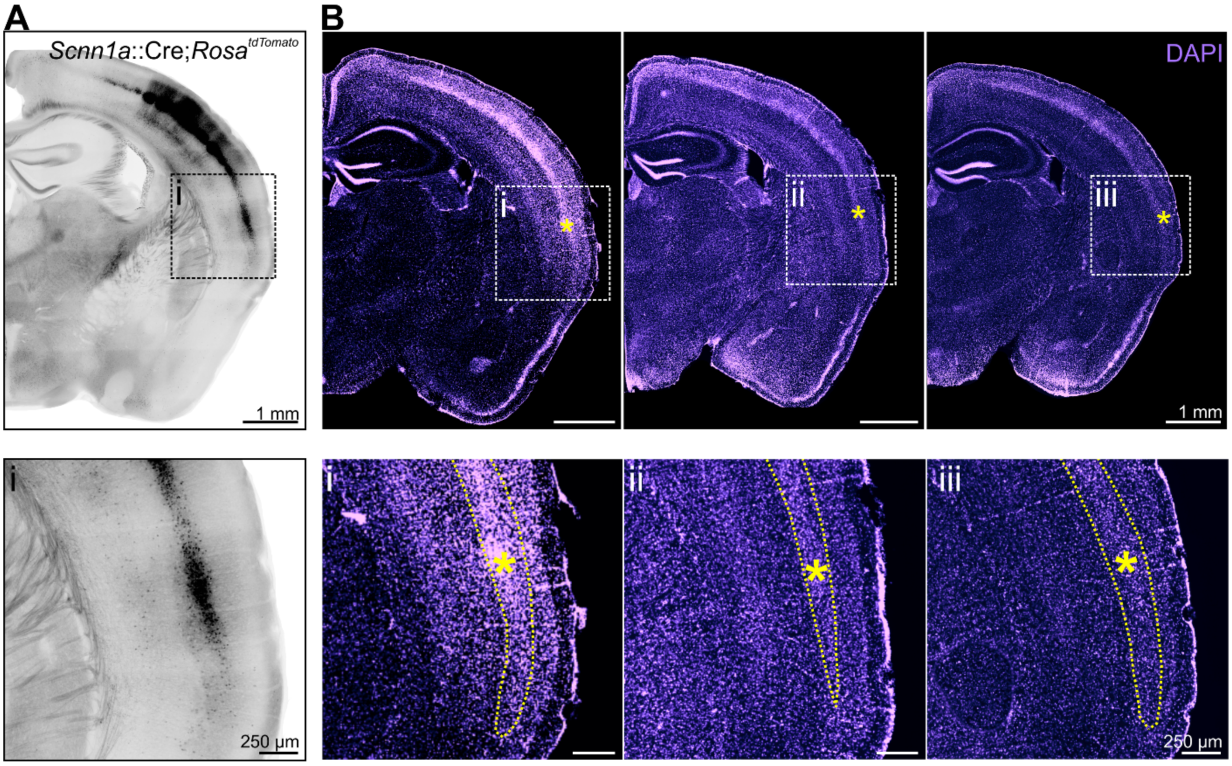


**Supplementary Figure 2. Thermal pIC contains a granular layer**

**A**, Representative micrograph of a coronal brain slice from mouse line expressing tdT in layer 4 neurons (Scnn1a-tdTomato), with the thermal responsive region of pIC. Higher resolution image (below) shows a dense granular layer (black cell bodies) in the thermal region of interest (see also Supplementary Figure 1 of Vestergaard et al., 2022). **B**, Representative micrographs of DAPI staining from 3 example mice used in the pIC tracing dataset. Boxes show region with dense granular layer comparable to that seen in **Ai**. Yellow asterisk indicates the center of injection site, yellow dashed line represents border of granular layer 4.

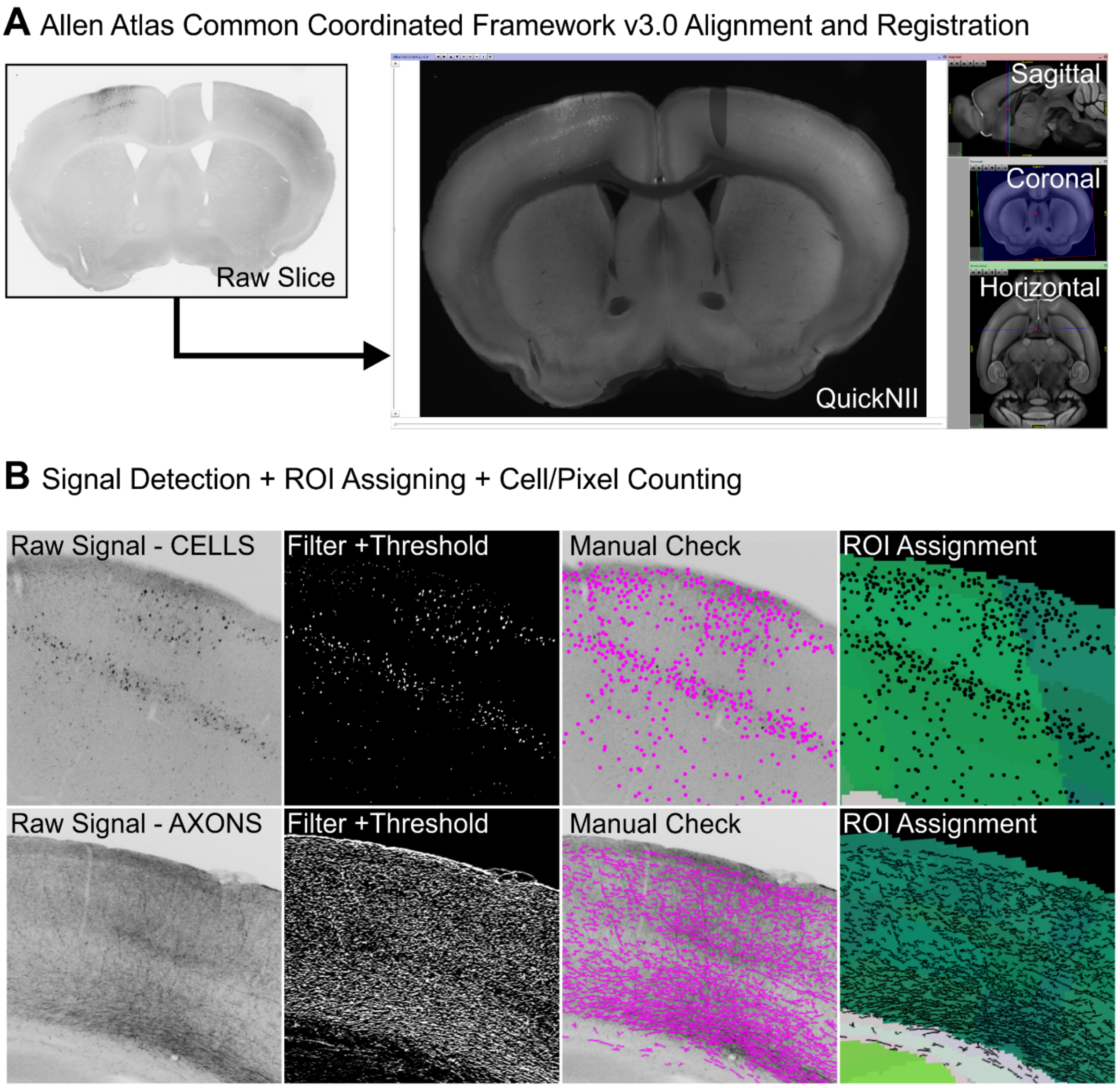


**Supplementary Figure 3. Analysis sequence to quantify whole-brain input/outputs.**

**A**, Brain sections were aligned and registered to the Allen Atlas Common Coordinated Framework v3.0 (Wang et al., 2020) using the QUICKNii Software (see Puchades et al., 2019). **B**, Top, signal filtering, thresholding, manual verification and ROI assignment of single cells projecting to either S1 or pIC. Bottom, same as top except for axons exiting S1 or pIC. Both examples are inputs to- or outputs from pIC.

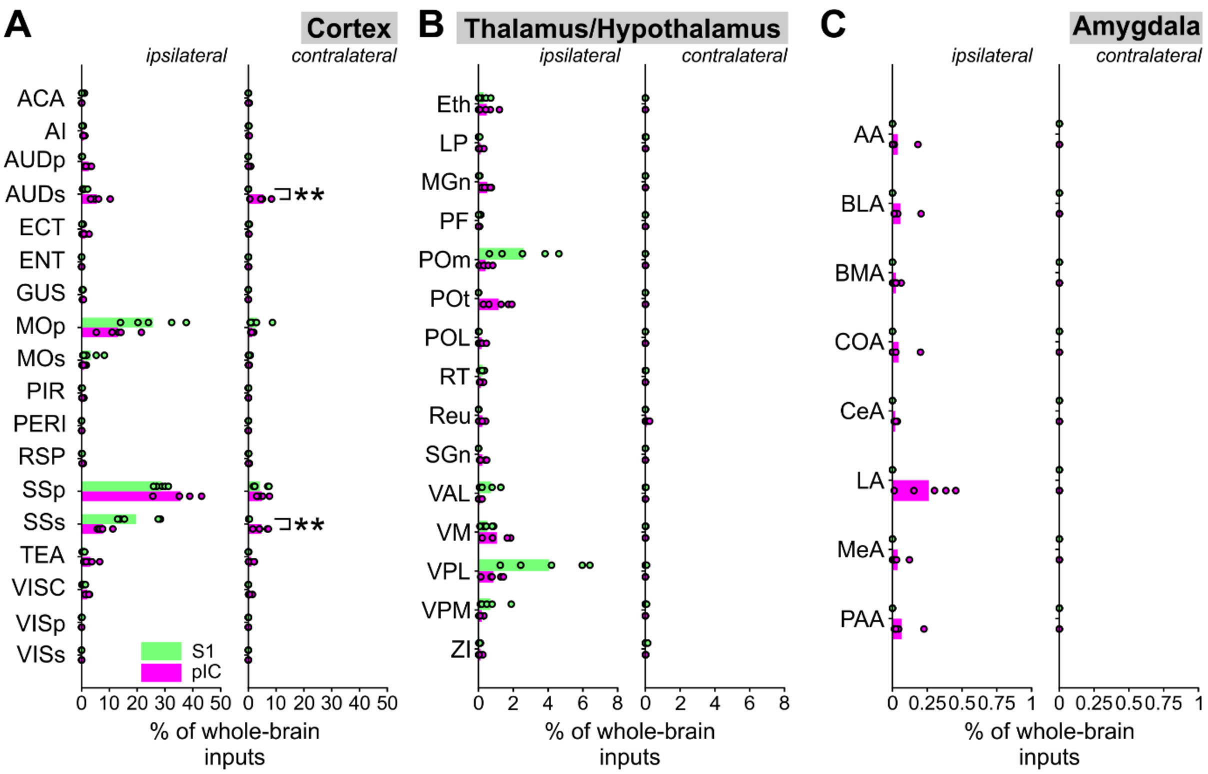


**Supplementary Figure 4. Ipsilateral and Contralateral inputs to thermal cortices.**

Comparison of ipsilateral (left) or contralateral (right) (**A**) cortical, (**B**) thalamic/hypothalamic, and (**C**) amygdaloid inputs to S1 (green) and pIC (magenta). Bars show means and open circles show individual mice, n = 5 mice per condition. * = $p < 0.05$, ** = $p < 0.01$, *** = $p < 0.001$. For detailed *p* values, see Supplementary Table 2. A full list of abbreviations is provided in Supplementary Table 1.

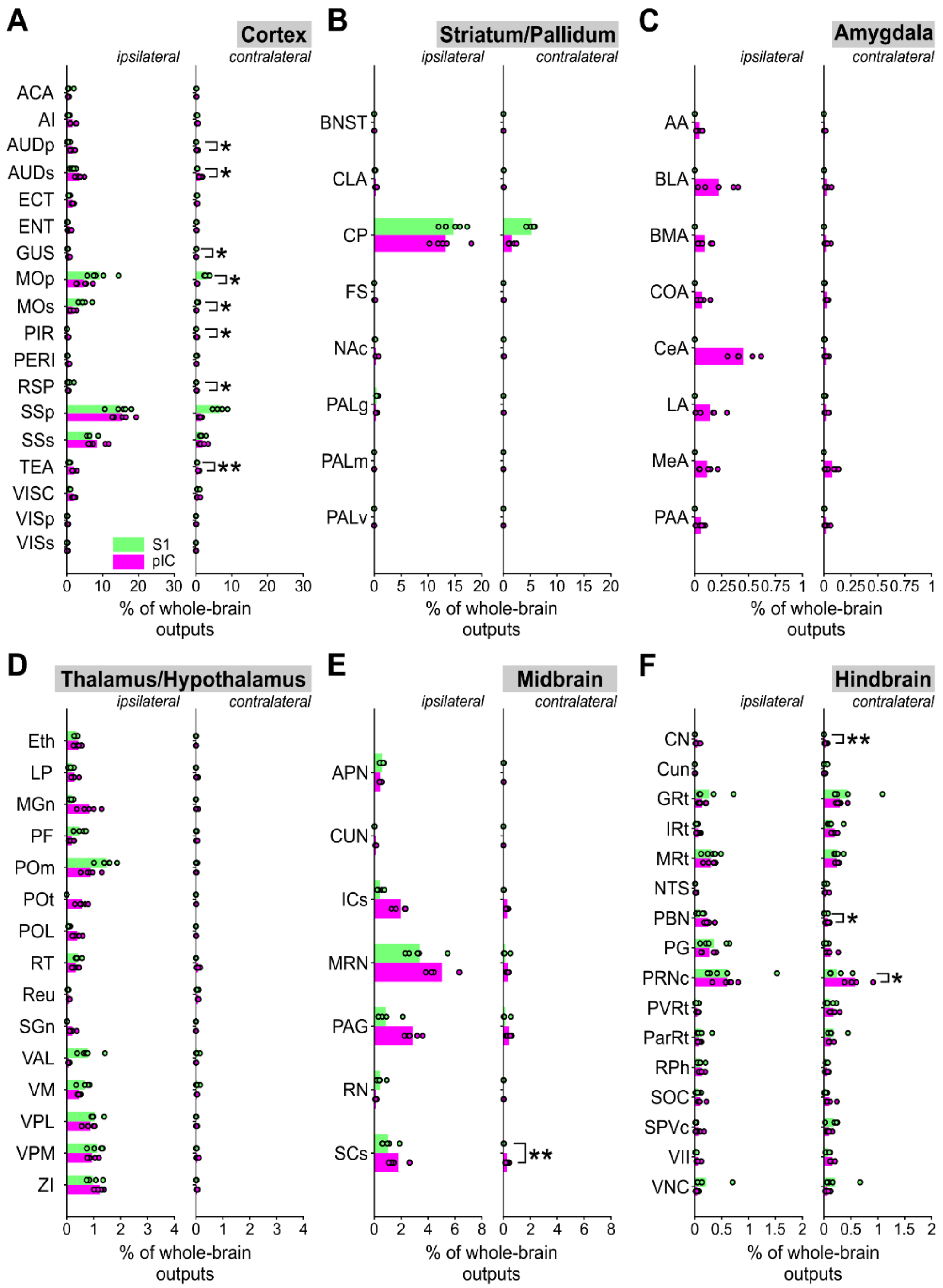


**Supplementary Figure 5. Ipsilateral and Contralateral outputs from thermal cortices.**

Comparison of S1 (green) and pIC (magenta) ipsilateral (left) or contralateral (right) outputs to subregions of the (**A**) cortex, (**B**) striatum/pallidum, (**C**) amygdala, (**D**) thalamus/hypothalamus, (**E**) midbrain, (**F**) hindbrain. Bars show means and open circles show individual mice, n = 5 mice per condition. * = $p < 0.05$, ** = $p < 0.01$, *** = $p < 0.001$. For detailed *p* values, see Supplementary Table 3. A full list of abbreviations is provided in Supplementary Table 1.

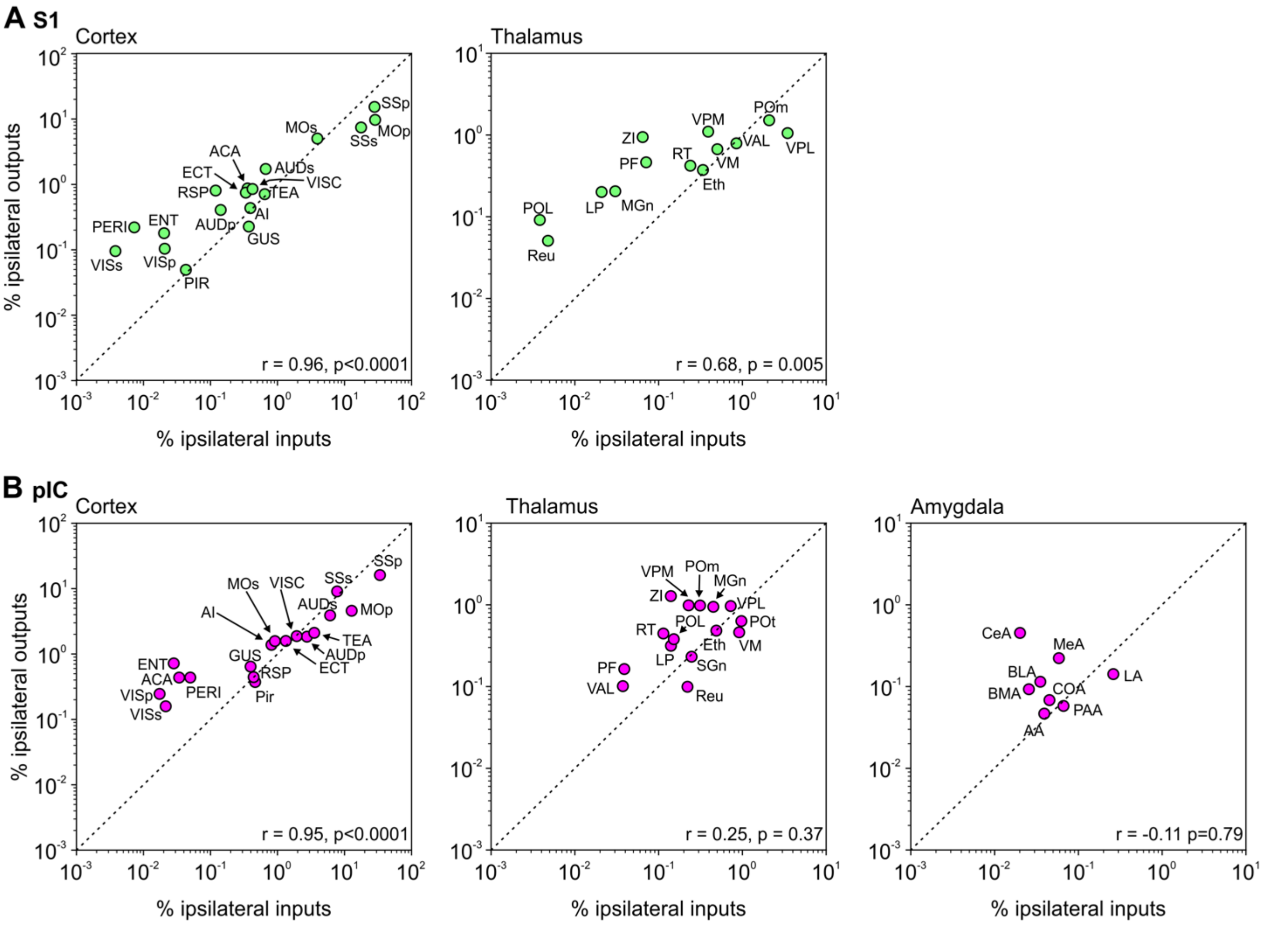


**Supplementary Figure 6. Correlations of input-output strength for S1 and pIC.**

**A**, Correlations of input-output strength for cortical and thalamic subregions connected to S1. Individual data points show mean value. r = Pearson's correlation coefficient. A full list of abbreviations is provided in Supplementary Table 1. **B**, Same as A. but for pIC and including connectivity with Amygdala.

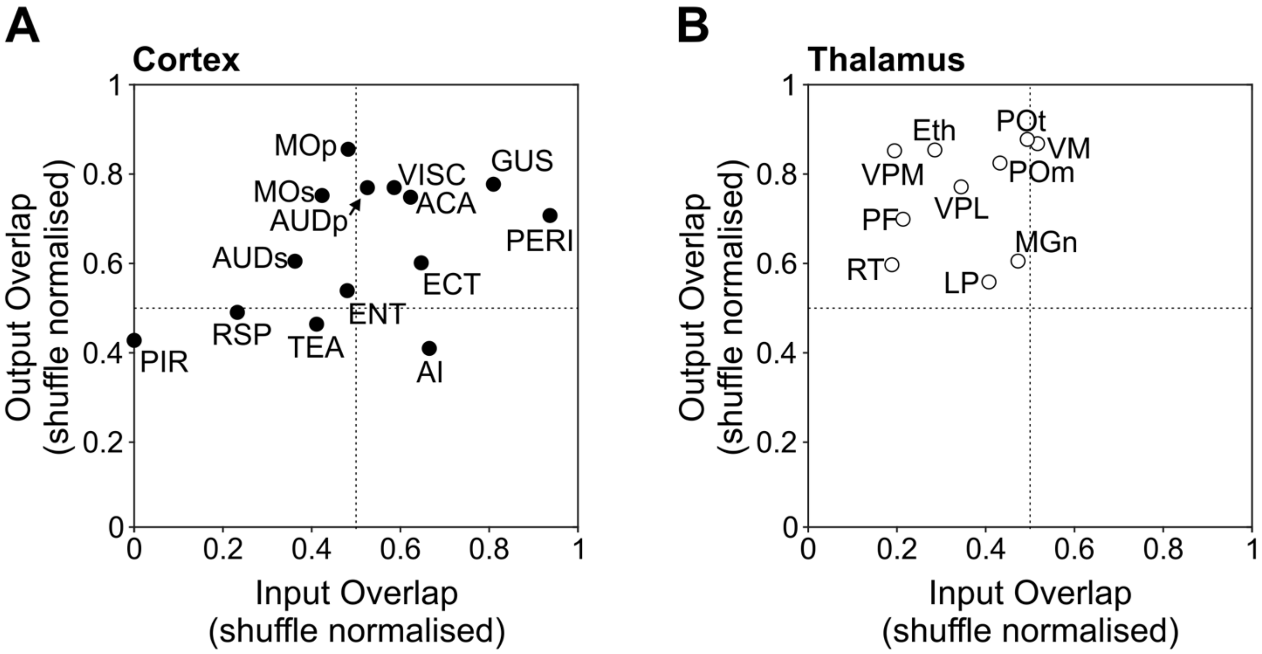


**Supplementary Figure 7. Input and output overlap coefficients normalized by the mean of the shuffled distribution.**

**A**, Input and output overlap coefficients normalized by the shuffle distribution mean for each cortical region. The position of the data points indicate similar amounts of input/output spatial overlap within cortical regions. Each closed circle denotes an individual brain region. A list of abbreviations is provided in Supplementary Table 1. **B**, Same as **A**, except for thalamic sub regions, with the position of data points encompassing the upper left quadrant, indicating that there is a greater different in their input/output spatial overlap . A list of abbreviations is provided in Supplementary Table 1. Each open circle denotes an individual brain region. A list of abbreviations is provided in Supplementary Table 1.

### Supplementary Table 1 – Abbreviation List

| **Brain Region** | **Abbreviation** |
|---|---|
| Anterior amygdalar | AA |
| Anterior cingulate area | ACA |
| Agranular insular | AI |
| Anterior pretectal nucleus | APN |
| Primary auditory | AUDp |
| Dorsal auditory area | AUDs |
| Posterior auditory area | AUDs |
| Ventral auditory area | AUDs |
| Basolateral amygdalar nucleus | BLA |
| Basomedial amygdalar | BMA |
| Bed nuclei of the stria terminalis | BNST |
| Central amygdalar | CeA |
| Claustrum | CLA |
| Dorsal cochlear | CN |
| Ventral cochlear | CN |
| Cortical amygdalar | COA |
| Caudoputamen | CP |
| Cuneate nucleus | Cun |
| Cuneiform nucleus | CUN |
| Extended amygdalar | EA |
| Ectorhinal | ECT |
| Entorhinal | ENT |
| Ethmoid nucleus of the thalamus | Eth |
| Fundus of striatum | FS |
| Gigantocellular | GRt |

| | |
|---|---|
| Gustatory area | GUS |
| Inferior colliculus | ICs |
| Intermediate reticular | IRt |
| Lateral amygdalar | LA |
| Lateral posterior | LP |
| Medial amygdalar | MeA |
| Medial geniculate | MGn |
| Primary motor | MOp |
| Secondary motor | MOs |
| Midbrain reticular nucleus | MRN |
| Magnocellular reticular | MRt |
| Nucleus accumbens | NAc |
| Nucleus of the solitary tract | NTS |
| Piriform-amygdalar area | PAA |
| Periaqueductal gray | PAG |
| Globus pallidus | PALg |
| Medial pallidum | PALm |
| Vental pallidum | PALv |
| Parietal | PAR |
| Paragigantocellular reticular | ParRt |
| Parabrachial nucleus | PBN |
| Perirhinal | PERI |
| Parafascicular nucleus | PF |
| Pontine gray | PG |
| Piriform area | PIR |
| Posterior limiting | POL |
| Posterior complex | POm |

| | |
|---|---|
| Posterior triangular | POt |
| Pontine reticular nucleus | PRNc |
| Parvicellular reticular | PVRt |
| Central linear nucleus raphe | RAmb |
| Dorsal nucleus raphe | RAmb |
| Rostral linear nucleus raphe | RAmb |
| Nucleus of reuniens | Reu |
| Red nucleus | RN |
| Nucleus raphe | RPh |
| Retrosplenial | RSP |
| Reticular nucleus of the thalamus | RT |
| Superior colliculus | SCs |
| Suprageniculate | SGn |
| Substantia nigra | SNr |
| Superior olivary complex | SOC |
| Spinal nucleus of the trigeminal | SPVc |
| Primary somatosensory | SSp |
| Supplemental somatosensory | SSs |
| Temporal association | TEA |
| Ventral anterior-lateral | VAL |
| Facial nucleus | VII |
| Visceral area | VISC |
| Primary visual area | VISp |
| Anteromedial visual | VISs |
| Ventral medial nucleus | VM |
| Lateral vestibular nucleus | VNC |
| Medial vestibular nucleus | VNC |

| | |
|---|---|
| Ventral posterolateral | VPL |
| Ventral posteromedial | VPM |
| Zona incerta | ZI |

**Supplementary Table 2 – Whole brain inputs**

| | pIC ipsi mean (%) | pIC ipsi SEM (%) | pIC contra mean (%) | pIC contra SEM (%) | S1 ipsi mean (%) | S1 ipsi SEM (%) | S1 contra mean (%) | S1 contra SEM (%) | ipsi t value | ipsi p value | contra t value | contra p value |
|---|---|---|---|---|---|---|---|---|---|---|---|---|
| **AI** | 0.8 | 0.13 | 0.21 | 0.05 | 0.35 | 0.09 | 0.13 | 0.04 | -2.9109 | 0.0196 | -1.3419 | 0.2165 |
| **AUDp** | 2.5 | 0.51 | 0.23 | 0.16 | 0.16 | 0.05 | | | -4.5613 | 0.0018 | -1.485 | 0.1758 |
| **AUDs** | 5.54 | 1.31 | 5.36 | 1.45 | 0.97 | 0.35 | 0.03 | 0.01 | -3.3598 | 0.0099 | -3.6593 | 0.0064 |
| **CIN** | 0.03 | 0.01 | 0.11 | 0.07 | 0.29 | 0.2 | 0.03 | 0.02 | 1.2962 | 0.2311 | -1.1883 | 0.2688 |
| **ECT** | 1.22 | 0.43 | 0.16 | 0.02 | 0.28 | 0.09 | 0.19 | 0.07 | -2.1441 | 0.0644 | 0.5501 | 0.5973 |
| **ENT** | 0.02 | 0.02 | 0.07 | 0.03 | 0.02 | 0.01 | 0.04 | 0.01 | 0.0239 | 0.9815 | -1.0087 | 0.3427 |
| **GUS** | 0.45 | 0.07 | 0.01 | 0 | 0.37 | 0.05 | 0.03 | 0.01 | -0.917 | 0.3859 | 1.9606 | 0.0856 |
| **MOp** | 13.06 | 2.63 | 1.31 | 0.17 | 25.71 | 4.22 | 2.93 | 1.5 | 2.5444 | 0.0345 | 1.0772 | 0.3128 |
| **MOs** | 0.86 | 0.26 | 0.18 | 0.07 | 3.31 | 1.5 | 0.24 | 0.13 | 1.613 | 0.1454 | 0.4682 | 0.6522 |
| **PIR** | 0.4 | 0.16 | 0.05 | 0.01 | 0.08 | 0.04 | 0.04 | 0.02 | -1.9057 | 0.0931 | -0.3973 | 0.7016 |
| **Prhin** | 0.04 | 0.01 | 0.02 | 0.01 | 0.01 | 0.01 | 0.04 | 0.01 | -2.1555 | 0.0632 | 1.7087 | 0.1259 |

| | | | | | | | | | | | |
|---|---|---|---|---|---|---|---|---|---|---|---|
| **RSPLN** | 0.38 | 0.09 | 0.12 | 0.07 | 0.14 | 0.04 | 0.07 | 0.03 | -2.6503 | 0.0292 | -0.7092 | 0.4983 |
| **SSp** | 35.68 | 2.9 | 4.86 | 0.8 | 28.69 | 0.98 | 4.18 | 1.25 | -2.2782 | 0.0522 | -0.4546 | 0.6615 |
| **SSs** | 7.72 | 0.95 | 4.73 | 1.04 | 19.67 | 3.44 | 0.22 | 0.05 | 3.352 | 0.0101 | -4.3144 | 0.0026 |
| **TEA** | 3.15 | 0.99 | 1.06 | 0.46 | 0.74 | 0.2 | 0.04 | 0.03 | -2.3793 | 0.0446 | -2.2228 | 0.0569 |
| **VISC** | 2.07 | 0.29 | 0.47 | 0.26 | 0.62 | 0.24 | 0.01 | 0.01 | -3.84 | 0.0049 | -1.7599 | 0.1165 |
| **VISp** | 0.02 | 0.01 | 0.01 | 0.01 | 0.05 | 0.03 | 0.01 | 0.01 | 1.0872 | 0.3086 | 0.0492 | 0.962 |
| **VISs** | 0.02 | 0.01 | 0.01 | 0.01 | 0.01 | 0.01 | 0 | 0 | -0.5818 | 0.5767 | 0 | 0 |
| **Eth** | 0.47 | 0.22 | 0 | 0 | 0.28 | 0.13 | 0 | 0 | -0.7665 | 0.4654 | 0 | 0 |
| **LP** | 0.12 | 0.05 | 0 | 0 | 0.03 | 0.01 | 0 | 0 | -1.5408 | 0.1619 | 0 | 0 |
| **MGn** | 0.5 | 0.1 | 0 | 0 | 0.03 | 0.01 | 0 | 0 | -4.4697 | 0.0021 | 0 | 0 |
| **PF** | 0.03 | 0.01 | 0 | 0 | 0.06 | 0.03 | 0 | 0 | 0.8533 | 0.4183 | 0 | 0 |
| **POm** | 0.41 | 0.13 | 0 | 0 | 2.59 | 0.74 | 0 | 0 | 2.8866 | 0.0203 | 0 | 0 |
| **POt** | 1.16 | 0.32 | 0 | 0 | 0 | 0 | 0 | 0 | -3.6702 | 0.0063 | 0 | 0 |
| **PiI** | 0.18 | 0.08 | 0 | 0 | 0.01 | 0.01 | 0 | 0 | -2.1547 | 0.0633 | 0 | 0 |
| **RT** | 0.14 | 0.04 | 0 | 0 | 0.23 | 0.05 | 0 | 0 | 1.4621 | 0.1819 | 0 | 0 |
| **Reu** | 0.22 | 0.06 | 0 | 0 | 0 | 0 | 0 | 0 | -3.3623 | 0.0099 | 0 | 0 |
| **SGn** | 0.22 | 0.08 | 0 | 0 | 0 | 0 | 0 | 0 | -2.591 | 0.0321 | 0 | 0 |

| | | | | | | | | | | | | |
|---|---|---|---|---|---|---|---|---|---|---|---|---|
| **VAL** | 0.07 | 0.04 | 0 | 0 | 0.72 | 0.26 | 0 | 0 | 2.4725 | 0.0386 | 0 | 0 |
| **VM** | 1.06 | 0.3 | 0 | 0 | 0.49 | 0.15 | 0 | 0 | -1.6932 | 0.1289 | 0 | 0 |
| **VPL** | 0.87 | 0.23 | 0 | 0 | 4.05 | 0.99 | 0 | 0 | 3.1235 | 0.0142 | 0 | 0 |
| **VPM** | 0.2 | 0.06 | 0 | 0 | 0.69 | 0.32 | 0 | 0 | 1.5022 | 0.1715 | 0 | 0 |
| **ZI** | 0.12 | 0.05 | 0 | 0 | 0.07 | 0.02 | 0 | 0 | -1.0458 | 0.3262 | 0 | 0 |
| **AA** | 0.04 | 0.04 | 0 | 0 | 0 | 0 | 0 | 0 | -1.0849 | 0.3096 | 0 | 0 |
| **BLA** | 0.06 | 0.04 | 0 | 0 | 0 | 0 | 0 | 0 | -1.5818 | 0.1523 | 0 | 0 |
| **BMA** | 0.03 | 0.01 | 0 | 0 | 0 | 0 | 0 | 0 | -2.4149 | 0.0422 | 0 | 0 |
| **CORT** | 0.05 | 0.04 | 0 | 0 | 0 | 0 | 0 | 0 | -1.148 | 0.2841 | 0 | 0 |
| **CeA** | 0.02 | 0 | 0 | 0 | 0 | 0 | 0 | 0 | -4.7309 | 0.0015 | 0 | 0 |
| **LA** | 0.26 | 0.08 | 0 | 0 | 0 | 0 | 0 | 0 | -3.2693 | 0.0114 | 0 | 0 |
| **MeA** | 0.04 | 0.02 | 0 | 0 | 0 | 0 | 0 | 0 | -1.5792 | 0.1529 | 0 | 0 |
| **PirA** | 0.07 | 0.04 | 0 | 0 | 0 | 0 | 0 | 0 | -1.6477 | 0.138 | 0 | 0 |

**Supplementary Table 3 - Whole brain outputs**

| | pIC ipsi mean (%) | pIC ipsi SEM (%) | pIC contra mean (%) | pIC contra SEM (%) | S1 ipsi mean (%) | S1 ipsi SEM (%) | S1 contra mean (%) | S1 contra SEM (%) | ipsi t value | ipsi p value | contra t value | contra p value |
|---|---|---|---|---|---|---|---|---|---|---|---|---|
| **AI** | 1.65 | 0.39 | 0.34 | 0.07 | 0.45 | 0.12 | 0.22 | 0.05 | -2.91 | 0.02 | -1.4406 | 0.1877 |
| **AUDp** | 1.65 | 0.29 | 0.28 | 0.09 | 0.4 | 0.15 | 0.06 | 0.02 | -3.88 | 0.005 | -2.372 | 0.0451 |
| **AUDs** | 3.57 | 0.44 | 1.12 | 0.23 | 1.76 | 0.3 | 0.36 | 0.05 | -3.36 | 0.01 | -3.2244 | 0.0122 |
| **CIN** | 0.42 | 0.06 | 0.12 | 0.03 | 0.77 | 0.31 | 0.1 | 0.02 | 1.08 | 0.312 | -0.5643 | 0.588 |
| **ECT** | 1.6 | 0.13 | 0.32 | 0.06 | 0.7 | 0.09 | 0.28 | 0.04 | -5.79 | 4E-04 | -0.5719 | 0.5831 |
| **ENT** | 0.84 | 0.24 | 0.13 | 0.03 | 0.18 | 0.05 | 0.11 | 0.04 | -2.73 | 0.026 | -0.326 | 0.7528 |
| **GUS** | 0.69 | 0.11 | 0.07 | 0.01 | 0.22 | 0.05 | 0.12 | 0.02 | -4.02 | 0.004 | 2.5106 | 0.0363 |
| **MOp** | 4.69 | 0.86 | 0.34 | 0.04 | 9.22 | 1.5 | 2.95 | 0.31 | 2.625 | 0.03 | 8.4356 | 0 |
| **MOs** | 1.65 | 0.36 | 0.11 | 0.02 | 4.64 | 0.72 | 0.35 | 0.09 | 3.71 | 0.006 | 2.6045 | 0.0314 |
| **PIR** | 0.39 | 0.06 | 0.19 | 0.05 | 0.06 | 0.02 | 0.04 | 0.01 | -5.42 | 6E-04 | -2.7434 | 0.0253 |
| **Prhin** | 0.5 | 0.1 | 0.13 | 0.02 | 0.22 | 0.04 | 0.16 | 0.05 | -2.72 | 0.026 | 0.4671 | 0.6529 |

| | | | | | | | | | | | |
|---|---|---|---|---|---|---|---|---|---|---|---|
| **RSPLN** | 0.41 | 0.06 | 0.19 | 0.04 | 0.69 | 0.36 | 0.08 | 0.02 | 0.755 | 0.472 | -2.4411 | 0.0405 |
| **SSp** | 15.44 | 1.21 | 1.39 | 0.17 | 15.04 | 1.26 | 6.81 | 0.71 | -0.23 | 0.827 | 7.3809 | 0.0001 |
| **SSs** | 8.46 | 1.16 | 1.85 | 0.44 | 7.18 | 0.67 | 1.7 | 0.31 | -0.96 | 0.367 | -0.2876 | 0.7809 |
| **TEA** | 2 | 0.23 | 0.69 | 0.09 | 0.71 | 0.1 | 0.28 | 0.03 | -5.13 | 9E-04 | -4.2625 | 0.0028 |
| **VISC** | 1.9 | 0.16 | 0.59 | 0.17 | 0.89 | 0.08 | 0.55 | 0.16 | -5.66 | 5E-04 | -0.1593 | 0.8774 |
| **VISp** | 0.23 | 0.05 | 0.06 | 0.02 | 0.1 | 0.05 | 0.02 | 0.01 | -2 | 0.081 | -1.9238 | 0.0906 |
| **VISs** | 0.13 | 0.06 | 0.02 | 0.01 | 0.08 | 0.05 | 0 | 0 | -0.74 | 0.479 | -0.9935 | 0.3496 |
| **BNST** | 0.02 | 0.01 | 0.01 | 0.01 | 0 | 0 | 0 | 0 | -2.48 | 0.038 | -1.6972 | 0.1281 |
| **CLA** | 0.32 | 0.08 | 0.06 | 0.01 | 0.11 | 0.04 | 0.06 | 0.01 | -2.37 | 0.045 | -0.4631 | 0.6557 |
| **CP** | 13.27 | 1.31 | 1.51 | 0.29 | 14.72 | 0.94 | 5.25 | 0.28 | 0.901 | 0.394 | 9.1591 | 0 |
| **FS** | 0.09 | 0.03 | 0.04 | 0.02 | 0.01 | 0.01 | 0.01 | 0 | -2.34 | 0.047 | -1.6836 | 0.1308 |
| **NAc** | 0.38 | 0.11 | 0.07 | 0.03 | 0.12 | 0.03 | 0.03 | 0.01 | -2.22 | 0.057 | -1.4979 | 0.1725 |
| **PALg** | 0.38 | 0.05 | 0.05 | 0.01 | 0.49 | 0.07 | 0 | 0 | 1.293 | 0.232 | -3.5791 | 0.0072 |
| **PALm** | 0 | 0 | 0 | 0 | 0 | 0 | 0 | 0 | 0 | 0 | 0 | 0 |
| **PALv** | | 0 | 0 | 0 | 0 | 0 | 0 | 0 | 0 | 0 | 0 | 0 |
| **Eth** | 0.44 | 0.06 | 0 | 0 | 0.35 | 0.03 | 0 | 0 | -1.31 | 0.226 | 0.098 | 0.9244 |
| **LP** | 0.29 | 0.05 | 0.03 | 0.02 | 0.19 | 0.04 | 0 | 0 | -1.51 | 0.169 | -1.8212 | 0.1061 |

| | | | | | | | | | | | |
|---|---|---|---|---|---|---|---|---|---|---|---|
| **MGn** | 0.83 | 0.16 | 0.02 | 0.02 | 0.2 | 0.03 | 0 | 0 | -4 | 0.004 | -1.3642 | 0.2097 |
| **PF** | 0.18 | 0.04 | 0.02 | 0.01 | 0.47 | 0.1 | 0.01 | 0.01 | 2.742 | 0.025 | -0.5064 | 0.6262 |
| **POm** | 0.89 | 0.13 | 0.01 | 0.01 | 1.49 | 0.14 | 0.02 | 0.01 | 3.167 | 0.013 | 1.1091 | 0.2996 |
| **POt** | 0.57 | 0.08 | 0.01 | 0 | 0 | 0 | 0 | 0 | -6.81 | 1E-04 | -5.3403 | 0.0007 |
| **PiI** | 0.4 | 0.07 | 0.01 | 0 | 0.08 | 0.01 | 0 | 0 | -4.5 | 0.002 | -4.3268 | 0.0025 |
| **RT** | 0.34 | 0.05 | 0.08 | 0.02 | 0.41 | 0.04 | 0 | 0 | 1.039 | 0.329 | -3.5253 | 0.0078 |
| **Reu** | 0.1 | 0.01 | 0.08 | 0.01 | 0.05 | 0.01 | 0.05 | 0.02 | -4.75 | 0.001 | -1.8083 | 0.1082 |
| **SGn** | 0.21 | 0.05 | 0 | 0 | 0.01 | 0 | 0 | 0 | -4.14 | 0.003 | -0.7163 | 0.4942 |
| **VAL** | 0.09 | 0.01 | 0 | 0 | 0.79 | 0.17 | 0.05 | 0.03 | 4.061 | 0.004 | 1.6979 | 0.128 |
| **VM** | 0.46 | 0.02 | 0.03 | 0 | 0.67 | 0.09 | 0.06 | 0.03 | 2.385 | 0.044 | 0.7892 | 0.4527 |
| **VPL** | 0.88 | 0.09 | 0.02 | 0.01 | 1.04 | 0.09 | 0.01 | | 1.165 | 0.278 | -2.168 | 0.062 |
| **VPM** | 0.94 | 0.08 | 0.06 | 0.02 | 1.14 | 0.11 | 0.01 | 0.01 | 1.465 | 0.181 | -2.4914 | 0.0374 |
| **ZI** | 1.23 | 0.06 | 0.04 | 0.01 | 0.97 | 0.11 | 0.01 | 0 | -1.99 | 0.082 | -3.6365 | 0.0066 |
| **AA** | 0.05 | 0.01 | 0.01 | 0 | 0 | 0 | 0 | 0 | -3.87 | 0.005 | -2.7004 | 0.0271 |
| **BLA** | 0.22 | 0.07 | 0.03 | 0.01 | 0 | 0 | 0 | 0 | -3.08 | 0.015 | -2.4822 | 0.038 |
| **BMA** | 0.09 | 0.03 | 0.03 | 0.01 | 0 | 0 | 0 | 0 | -3.42 | 0.009 | -2.4467 | 0.0401 |
| **CORT** | 0.07 | 0.02 | 0.03 | 0 | 0 | 0 | 0 | 0 | -3.14 | 0.014 | -6.7438 | 0.0001 |

| | | | | | | | | | | | | |
|---|---|---|---|---|---|---|---|---|---|---|---|---|
| **CeA** | 0.45 | 0.05 | 0.02 | 0.01 | 0 | 0 | 0 | 0 | -8.24 | 0 | -3.5368 | 0.0077 |
| **LA** | 0.14 | 0.05 | 0.03 | 0.01 | 0 | 0 | 0 | 0 | -2.74 | 0.025 | -3.2366 | 0.0119 |
| **MeA** | 0.11 | 0.03 | 0.08 | 0.02 | 0 | 0 | 0 | 0 | -3.5 | 0.008 | -3.2002 | 0.0126 |
| **PirA** | 0.06 | 0.02 | 0.02 | 0.01 | 0 | 0 | 0 | 0 | -3.84 | 0.005 | -2.2282 | 0.0565 |
| **APN** | 0.45 | 0.03 | 0.02 | 0 | 0.62 | 0.06 | 0.01 | 0.01 | 2.448 | 0.04 | -0.6197 | 0.5527 |
| **CUN** | 0.12 | 0.02 | 0.01 | 0 | 0.03 | 0.01 | | | -5.49 | 6E-04 | -4.5154 | 0.002 |
| **ICs** | 1.96 | 0.22 | 0.29 | 0.03 | 0.43 | 0.1 | 0.01 | 0.01 | -6.42 | 2E-04 | -8.0088 | 0 |
| **MRN** | 5.04 | 0.54 | 0.32 | 0.04 | 3.39 | 0.55 | 0.14 | 0.09 | -2.13 | 0.066 | -1.7921 | 0.1109 |
| **PAG** | 2.84 | 0.25 | 0.42 | 0.08 | 0.85 | 0.34 | 0.14 | 0.1 | -4.79 | 0.001 | -2.228 | 0.0565 |
| **RN** | 0.13 | 0.02 | 0.02 | 0.01 | 0.44 | 0.13 | 0.01 | 0.01 | 2.404 | 0.043 | -0.9834 | 0.3542 |
| **SCs** | 1.82 | 0.34 | 0.27 | 0.05 | 1.04 | 0.23 | 0.03 | 0.01 | -1.9 | 0.094 | -4.6291 | 0.0017 |
| **CN** | 0.05 | 0.02 | 0.03 | 0.01 | 0 | 0 | 0 | 0 | -2.7 | 0.027 | -3.4142 | 0.0092 |
| **Cun** | 0 | 0 | 0.01 | 0.01 | 0 | 0 | 0.01 | 0.01 | -0.97 | 0.362 | 0.5311 | 0.6098 |
| **GRt** | 0.13 | 0.03 | 0.3 | 0.04 | 0.27 | 0.12 | 0.44 | 0.17 | 1.082 | 0.311 | 0.7823 | 0.4565 |
| **IRt** | 0.07 | 0.01 | 0.2 | 0.02 | 0.03 | 0.01 | 0.16 | 0.05 | -2.17 | 0.062 | -0.7864 | 0.4543 |
| **MRt** | 0.3 | 0.04 | 0.24 | 0.02 | 0.31 | 0.06 | 0.25 | 0.03 | 0.107 | 0.917 | 0.1779 | 0.8632 |
| **NTS** | 0.02 | 0.01 | 0.04 | 0.02 | 0.01 | 0 | 0.02 | 0.01 | -2.2 | 0.059 | -0.6736 | 0.5196 |

| | | | | | | | | | | | | |
|---|---|---|---|---|---|---|---|---|---|---|---|---|
| **PBN** | 0.26 | 0.03 | 0.07 | 0.01 | 0.1 | 0.03 | 0.02 | 0.01 | -3.63 | 0.007 | -2.7913 | 0.0235 |
| **PG** | 0.27 | 0.06 | 0.12 | 0.04 | 0.36 | 0.11 | 0.04 | 0.01 | 0.705 | 0.501 | -1.9234 | 0.0906 |
| **PRNc** | 0.61 | 0.08 | 0.59 | 0.09 | 0.62 | 0.24 | 0.24 | 0.08 | 0.034 | 0.974 | -2.8074 | 0.0229 |
| **PVRt** | 0.05 | 0.01 | 0.18 | 0.03 | 0.03 | 0.01 | 0.11 | 0.04 | -1.4 | 0.199 | -1.2129 | 0.2598 |
| **ParRt** | 0.08 | 0.01 | 0.12 | 0.02 | 0.12 | 0.05 | 0.18 | 0.07 | 0.857 | 0.417 | 0.827 | 0.4322 |
| **RPh** | 0.13 | 0.02 | 0.06 | 0.01 | 0.11 | 0.03 | 0.05 | 0.01 | -0.7 | 0.503 | -0.7621 | 0.4679 |
| **SOC** | 0.11 | 0.03 | 0.1 | 0.04 | 0.05 | 0.02 | 0.02 | 0.01 | -1.61 | 0.146 | -1.9514 | 0.0868 |
| **SPVc** | 0.07 | 0.03 | 0.09 | 0.02 | 0.02 | 0.01 | 0.18 | 0.04 | -1.61 | 0.147 | 1.9962 | 0.081 |
| **VII** | 0.06 | 0.02 | 0.16 | 0.02 | 0.02 | 0.01 | 0.08 | 0.02 | -2.29 | 0.051 | -3.0886 | 0.0149 |
| **VNC** | 0.05 | 0.01 | 0.07 | 0.02 | 0.21 | 0.12 | 0.21 | 0.12 | 1.312 | 0.226 | 1.1869 | 0.2693 |

**Supplementary Movie 1 – Whole brain inputs**

**Supplementary Movie 2 – Whole brain outputs**
